# Genome-wide dynamic nascent transcript profiles reveal that most paused RNA polymerases terminate

**DOI:** 10.1101/2025.03.27.645809

**Authors:** Rudradeep Mukherjee, Michael J. Guertin

## Abstract

We present a simple model for analyzing and interpreting data from kinetic experiments that measure engaged RNA polymerase occupancy. The framework represents the densities of nascent transcripts within the pause region and the gene body as steady-state values determined by four key transcriptional processes: initiation, pause release, premature termination, and elongation. We validate the model’s predictions using data from experiments that rapidly inhibit initiation and pause release. The model successfully classified factors based on the steps in early transcription that they regulate, confirming TBP and ZNF143 as initiation factors and HSF and GR as pause release factors. We found that most paused polymerases terminate and paused polymerases are short-lived with half lives less than a minute. We make this model available as software to serve as a quantitative tool for determining the kinetic mechanisms of transcriptional regulation.

## Introduction

Gene expression regulation plays a crucial role in determining cell fate during development and influences how cells respond to environmental signals. Early events in the transcription cycle are major determinants of a gene’s transcriptional output. The earliest events of transcription include the formation of a preinitiation complex, promoter opening, rounds of abortive initiation, promoter escape, and ultimately pausing of RNA polymerases 50–100 bases down-stream from the transcription start site (1–3). We refer to these early events collectively as *initiation* throughout this study because most molecular measurements in cells cannot distinguish between these transient and unstable states. After initiation, the paused RNA polymerases may either proceed into productive elongation or prematurely terminate and disassociate from the DNA (4–6). Regulation of these steps is crucial for the precise control of transcription.

Previous studies have used imaging or high-throughput genomics assays to investigate and model the kinetics of transcription. Imaging studies track the generation of transcripts linked to fluorescent reporters and use models to interpret the time series of fluorescence signal (7, 8). Others use fluorescence recovery after photobleaching or photoactivatable molecules to determine residence times of RNA polymerases in different stages of transcription (9, 10). Genome-wide nuclear run-on assays, which precisely measure the position and orientation of transcriptionally engaged RNA polymerases, have been combined with inhibitors of initiation and pause release to dissect contributions of each regulated step (11). A recent study employed generative probabilistic modeling of nascent transcription datasets to estimate rates of initiation and pause release (12), highlighting the rate-limiting nature of RNA polymerase escape from the pause region. Despite the numerous studies and diverse methodologies employed, there is no consensus on the absolute and relative rates of these key regulatory steps.

Transcription factors (TFs) interact with various co-factors to coordinate the recruitment and activity of RNA polymerases (13–25). TFs bind to sequence motifs and assemble complexes that facilitate the loading and progression of RNA polymerases (26–29). Understanding the molecular functions of transcription factors, specifically how they contribute to the regulation of RNA polymerase activity at these various steps in early elongation, is crucial to understanding the complex mechanisms that drive gene expression (30, 31). Many foundational studies have leveraged rapidly inducible transcription factors to reveal their regulatory roles by tracking changes in RNA polymerase profiles at target genes (31–36). New chemical and physical methods can more broadly perturb TF activity or degrade TFs, significantly expanding the range of factors whose molecular functions can be studied (37–42). Determining the specific role of each transcription factor and their combinatorial effects on transcription regulation requires methods to acutely activate or perturb TF activity, quantify changes in RNA polymerase activity, and a modeling framework to interpret the data.

We present a Compartment Model framework, which represents RNA polymerase densities in the pause region and gene body as steady-state values determined by four key transcriptional processes: initiation, pause release, premature termination, and elongation (Fig. 1A). Unless stated otherwise, we define *initiation* as the sequence of events spanning from the incorporation of the first RNA base to the transition into the pause site. For each gene, the model estimates the magnitudes of pause release and initiation rates, as well as changes in these rates for all genes after experimental perturbation (Fig. 1B-C). We describe the framework and validate the model’s predictions through analysis of publicly available experimental data that inhibited either initiation or pause release. Analysis of triptolide-treated datasets, which inhibit initiation, reveals that premature termination occurs at a faster rate than pause release. We further demonstrate that pause release serves as a critical regulatory step that significantly influences transcription output, but only when the premature termination rate of paused polymerases exceeds the release rate of paused polymerases into the gene body. We analyze publicly available transcription factor activation and perturbation data to confirm that ZNF143 and TBP predominantly regulate initiation, while the glucocorticoid receptor (GR) and *Drosophila* heat shock factor (HSF) regulate pause release. The model is available as a command-line tool that processes relative pause and gene body RNA polymerase densities from two experimental conditions as inputs and provides estimates of initiation rates, pause release rates, premature termination rates, and absolute RNA polymerase densities in pause regions and gene bodies.

**Fig. 1.**
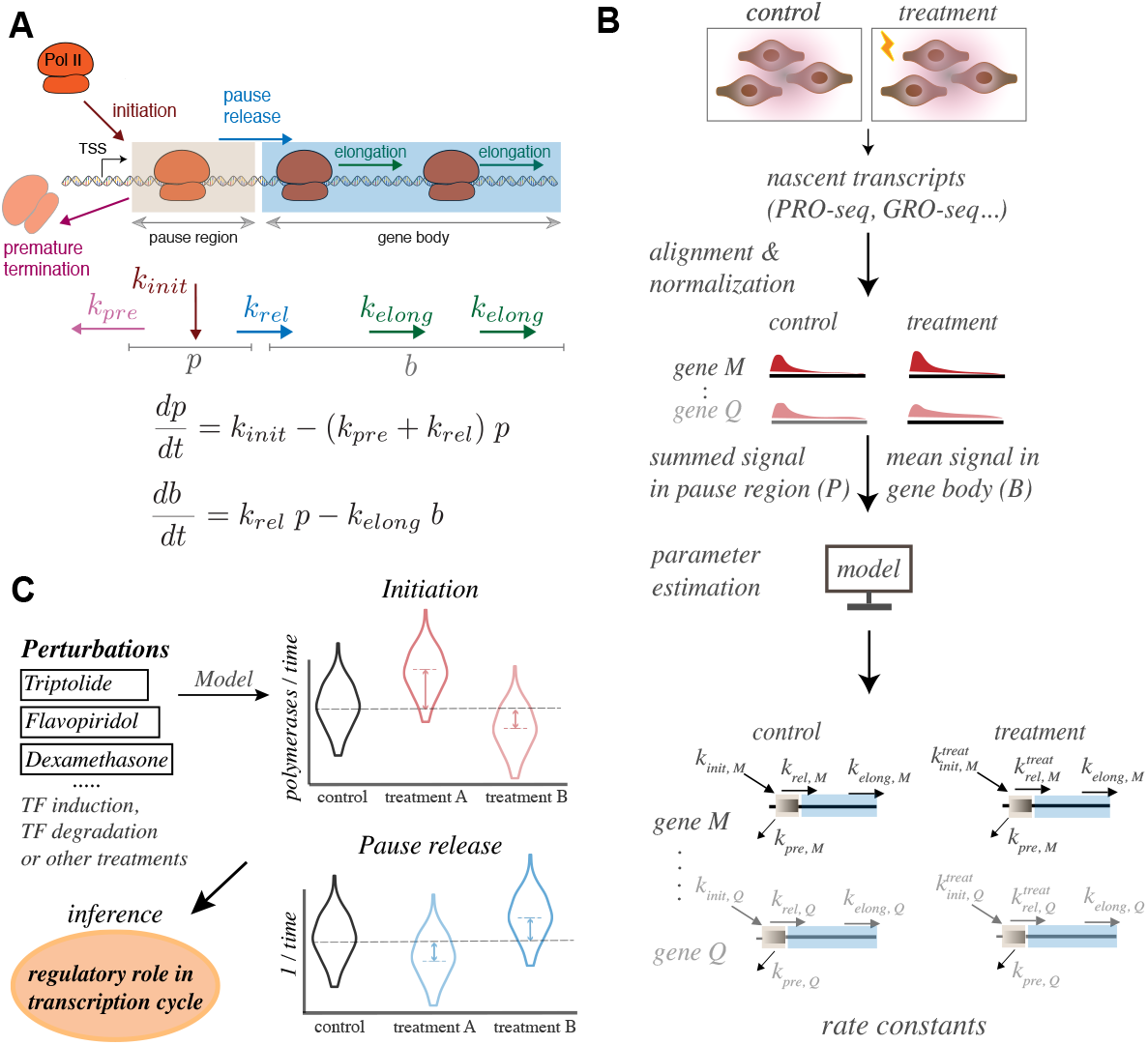
Kinetic modeling of transcription data determines the effect of treatments and perturbations. A) A gene has two compartments in the model–the pause window (*p*) and the gene body (*b*). RNA polymerase enters the pause region with an initiation rate constant *k*_*init*_, prematurely terminates with a rate constant *k*_*pre*_, escapes from pause region with a release rate *k*_*rel*_, and proceeds through the gene body based on the elongation rate *k*_*elong*_. Herein, we assume that *k*_*pre*_ and *k*_*elong*_ remain constant for a gene between conditions. B) The ideal input data for the model is generated from genome-wide nascent transcriptome (PRO-seq) experiments upon acute treatment with a perturbation or stimulus. Standard methods identify differentially expressed genes and the normalized data provides densities of reads in pause and gene body compartments. The model uses these densities to infer initiation and pause release rates for input genes. C) The regulatory effects on pause release and initiation, induced by general perturbations or the acute activation/inhibition of specific transcription factors, are determined by the rate changes explained by the model.

## Materials & Methods

### Processing of nascent transcript datasets

We processed six datasets spanning across a broad range of treatments: triptolide and flavopiridol in mouse v6.5 cells (11), degradation of TBP in HAP1 cells (43) and ZNF143 in HEK293T cells (44), dexamethasone treatment in A549, U2OS, C7 cells (45, 46), and heat shocked *Drosophila* S2 cells (33). We downloaded the datasets from Gene Expression Omnibus (Table S1) and analyzed the raw data using a general pipeline (47, 48). We normalized signal within datasets using `sizeFactors` from DESeq2 (49). Since triptolide (12.5 min) and flavopiridol (25 min) treatments cause universal repression, we normalized these datasets by using read counts from gene regions > 50 kb for triptolide (> 100 kb for flavopiridol) as *controlGenes* in DESeq2 (49). We chose these gene regions because an elongation rate of 4kb/min is beyond the higher end of contemporary estimates of elongation rates (50). We defined Heat Shock Factor (HSF) activated genes in S2 cells (33) by first aligning all data to an *Hsp70* consensus sequence. We mapped the remaining unmapped reads to the `dm6` genome, counted reads within genes, and normalized the replicates using DE-Seq2 `sizeFactors` (49). We only considered the 249 activated genes selected from Table S2 in Duarte et. al., (33) because these genes were carefully curated for read-through transcription. We considered 65 heat shock activated genes (FDR < 0.001) that are dependent upon the presence of HSF.

### Hsp70 consensus sequence

We generated the *Hsp70* consensus gene sequence by combining the *Hsp70* sequences from the BDGP6.46 (release 112) reference. We used the start and end coordinates of six Hsp70 genes (*Hsp70Aa/Hsp70Ab, Hsp70Ba, Hsp70Bb, Hsp70Bbb, Hsp70Bc*) and included an additional 50 bases upstream of the reference start site. Using these coordinates, we extracted the six Hsp70 sequences from the reference genome with `bedtools getfasta` (51). We then created an alignment file for these sequences using `clustalw` (52). The consensus sequence was derived by using Biopy-thon’s `dumb_consensus()` function (53) to the alignment file with a threshold of 0.6.

### Estimation of Pol II densities at pause region and gene body region

To define the pause region of a gene, we searched for a 50 bp window with the maximum summed PRO-seq signal within a 200 bp region (300 bp for GRO-seq) downstream from the most prominent transcription start site (TSS) as defined by `primaryTranscriptAnnotation` (54). This 50 bp window is designated the *pause region*. The pause density or pause sum is the summed signal across all base pairs in this window. We sum the reads in pause region to accommodate signals from RNA polymerase pausing at any arbitrary position within the pause window. We define the gene body as the region 500 bp from end of the pause window to the inferred transcription termination site (54) or the distance an initiated polymerase can travel during the time of treatment if RNA polymerase elongates at a rate of 2kb/min, whichever is shorter. For example, the body length of genes within the flavopiridol data set were constrained to 50 kb to capture the region where RNA polymerase clears out upon treatment. For genes repressed by 12.5 minutes of triptolide treatment, we quantified gene body density using a 4 kb window beginning 1 kb downstream of the pause region. This window was selected to minimize inclusion of the transitional gene-body region representing the period required for triptolide to reach 75% efficacy (55). We define the body density as the mean signal in this window.

### Steady-State Assumption for Pause Window and Gene Body Densities

We estimate pause and body densities from genomic nuclear run-on data for two conditions; we consider RNA polymerase densities to be at steady state for each condition. We validated these steady-state assumptions by using triptolide treatment and dexamethasone treatment datasets. We fit an exponential decay function to measured initiation rates following triptolide treatment (55). These data were generated using an *in vitro* transcription assay that quantified initiation from the CMV promoter (Fig. S1A). The slope of the exponential fit at 12.5 minutes was approximately −10−4 (Fig. S1B). This slope approaches the value of zero, which supports the assumption that initiation is at steady-state after 12.5 minutes of treatment.

To determine whether 45 minutes of dexamethasone treatment results in steady state regulation of dexamethasone-activated genes, we focused on genes longer than 150kb (45) (Fig. S1C-E). We analyzed long genes because the 3^′^ ends are not affected by the treatment and we expect a transition state that represents the non-steady state changes in density caused by dexamethasone. If the genes reach steady state, then the 5^′^ end of the gene will have a consistent RNA polymerase density leading into this transition state. We demarcated the gene bodies into three distinct regions: affected by dexamethasone, unaffected by dexamethasone, and a transition region (Fig. S1C). The relatively flat traces in affected and unaffected regions of dexamethasone-activated genes provide support that the non-steady state period is transient and short (Fig. S1D). We also selected 85 unchanged genes that were matched for both length and expression level (Fig. S1E). Therefore, the first 90kb of gene bodies can be considered approximately steady-state after 45 minutes of dexamethasone treatment.

### The Compartment Model

We describe the time evolution of pause and body densities in the following manner:

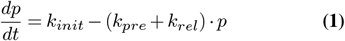

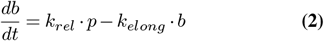

In the above equations, *t* is time, *p* is the density in pause region, *b* is mean density in gene body and *k*_*init*_, *k*_*rel*_, *k*_*pre*_, *k*_*elong*_ are rates of initiation, pause release, premature termination, and elongation, respectively. The measurements of pause sum and body densities from sequencing data are assumed to be steady-state values. The steady state assumption is reasonable because the measured rates of these various processes are much faster than the experimental treatment times. At steady-state, we have:

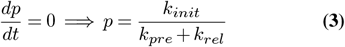

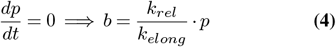

For each gene, we consider that initiation (*k*_*init*_) and pause release (*k*_*rel*_) rates are regulated between treatments, while elongation (*k*_*elong*_) and premature termination (*k*_*pre*_) rates remain constant. Although the range of elongation rates for genes tend to be 1.5-4 kb/min (11, 34, 56–58), we cannot accurately estimate average elongation rates for all genes for each condition. However, we make this assumption of constant elongation rate because RNA polymerase signal within gene bodies correlate well with accumulated RNA-seq signal (59). If elongation rate increased during gene activation, then RNA polymerase density would decrease as mRNA output increases.

Considering two conditions under study as *control* and *treatment*, we have the following relations for change in initiation and pause release rates. *FC* denotes fold change between treatment and control.

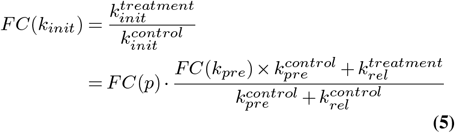

If premature termination for a gene does not change between treatments, we get:

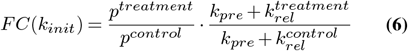

For change in pause release rate, we have:

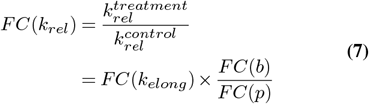

If we assume that elongation rate does not change for a gene between treatments, the change in pause release rate is simply the change in body density divided by the change in pause density:

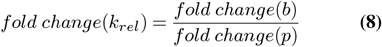

In Eq. 6, the fraction 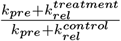 varies between two values depending upon relative values of premature termination rate (*k*_*pre*_) and pause release rate (*k*_*rel*_):

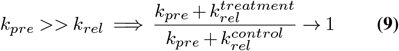

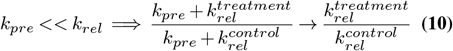

From Eq. 6, Eq. 9 and Eq. 10, fold change in initiation rate (*k*_*init*_) for a gene can vary between fold change in pause density and the fold change in body density:

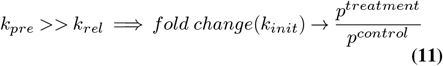

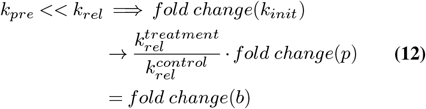

The fold change in pause release (Eq. 8) is an inverse ratio of change in *pausing index* (the ratio of pause density to body density). As accurate measures of genic elongation rate become available, we can calculate the change in pause release rate by dividing the change in elongation rate by the change in pausing index (Eq. 7). Note that the normalization or scaling of read signals to estimate steady-state *p* and *b* values does not affect the fold change in pause release (Eq. 8) and bounds on changes in initiation rate (Eqs. 11–12). Consequently, the inference of the regulatory effects of a perturbation on initiation and pause release does not depend on the scaling of pause and body densities. For absolute values of rates we derive the following from Eq. 3 & 4:

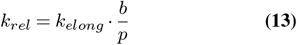

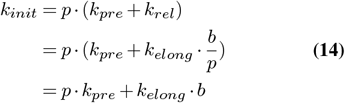

From Eq. 13, the absolute value of pause release rate (*k*_*rel*_) for a gene depends upon elongation rate (usually between 1800-4000 bp/min) and the inverse of the pause to body density ratio. Therefore, the value of pause release rate constant (*k*_*rel*_) is also not dependent upon normalization or scaling criteria. From Eq. 14, the initiation rate (*k*_*init*_) depends upon the absolute values of pause and body densities (i.e. the number of RNA polymerase in each compartment). Additionally, the amount of polymerases releasing into gene body, the effective release rate (*k*_*rel*_ × *p*), provides a way to calculate spacing between polymerases on the gene body in base pairs:

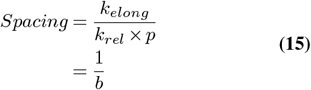

where, elongation rate (*k*_*elong*_) is in units of 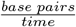 and pause release (*k*_*rel*_) has the same time unit. From Eq. 13, the magnitude of occupancy is also equal to reciprocal of body density (*b*).

We derive a relation between premature termination rate (*k*_*pre*_) and pause release rate (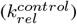) from Eq. 5 & Eq. 7:

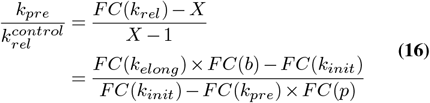

where 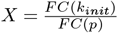 with *p* representing the pause density for a gene.

To ensure that the ratio in Eq. 16 remains positive, the fold change in initiation rate, *FC*(*k*_*init*_) should lie between two values - *FC*(*k*_*elong*_) × *FC*(*b*) and *FC*(*k*_*pre*_) × *FC*(*p*). If we assume that the rates of premature termination and elongation do not change for a gene between treatments, then the ratio of premature termination to pause release rate is a function of initiation fold change and fold change in the pause and body regions:

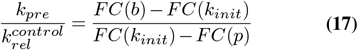

Given experimental data on the change in initiation rates after treatment, we assign the rates of premature termination to each gene promoter using the relations in Eqs. 16 & 17. Importantly, this ratio does not depend on the scaling criteria employed for pause and body densities, because the equations only consider their fold changes. The ratio in Eq. 17 becomes zero or undefined when the numerator or denominator becomes zero:

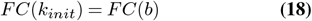

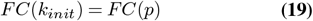

The values in Eqs. 18–19, which render 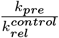 as zero or undefined, represent the extreme bounds on changes in initiation rate as described in Eqs. 11–12. However, transcription is expected to operate within a regime where the rates of premature termination (*k*_*pre*_) and pause release (*k*_*rel*_) yield a non-zero ratio for 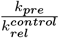. Consequently, the extreme cases for the relative values of *k*_*pre*_ and *k*_*rel*_, as described in Eqs. 11–12, are not considered.

The ratio of 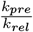 can be expressed using a variable that defines placement of *FC* (*k*_*init*_) within the interval defined by, *FC*(*p*) and *FC*(*b*). We consider, *FC*(*p*) < *FC*(*b*) without loss of generality, and assume *FC*(*k*_*init*_) = *FC*(*p*) + *Y* ∗ [*FC*(*b*) − *FC*(*p*)], where *Y* is a factor between 0 and 1:

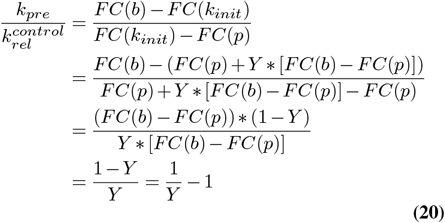

### Pause and body densities as occupancy values

To infer rates of initiation at different gene promoters we first transformed pause and gene body densities into absolute occupancy values. Occupancy refers to the number of RNA polymerases in the pause region or gene body per gene length. We assume that a fully occupied pause region will contain only one RNA Polymerase and that some genes would approach full occupancy of the pause region. In an ideal dataset without aneuploidy, gene duplications, PCR biases, and sequence mapping ambiguities, the ranked pause sums would saturate to a maximum occupancy. However, over all the datasets examined, the observed ranked pause sums exhibit a sudden uptick at the highest pause densities (Fig. S2). Manual inspection of genes with high pause signals revealed multicopy genes located within non-uniquely mappable regions and genes characterized by highly variable read densities in the gene body. We estimated a fully occupied pause region by fitting ranked pause densities of all expressed genes to a hyperbolic tangent function (*y* = *A* · *tanh*(*B* · *x*) + *C*) where density of a fully occupied pause region is assumed at *saturation* = *A* + *C*. We fitted the hyperbolic tangent function to log-transformed ranked pause sums using Scipy’s `optimize.curve_fit()` function (60). We evaluated both untransformed and log-transformed ranked pause sums and attempted to fit several saturation functions. Fitting log-transformed values with the hyperbolic tangent function provided consistent results across datasets with Unique Molecular Identifiers (UMIs). As we expand upon in the *Results*, our calculation of gene body densities with these values are consistent with imaging data (61–64).

We reasoned that a saturation value should be robust to amount of input data used to fit the curve, and be minimally influenced by the observed exponential increase in pause sums. For datasets with Unique Molecular Identifiers (UMIs) from which PCR duplicated reads could be confidently removed, the saturation values from incremental fits to pause densities (70th to 100th percentile) showed a modest increase by 1.5- to 4-fold (Fig. S2D-F & Fig. S3A). In contrast, the saturation values from incremental fits for datasets without UMIs varied from 23-fold to 600-fold (Fig. S2A-C & Fig. S3A). The saturation fits for datasets with UMIs tended to stabilize and reach 90th percentile of the ranked paused sums (Fig. S2D-F). Since the saturation values for datasets with UMIs were robust, we used the saturation values obtained from fits to all genes as the maximum occupancy signal. To remain consistent and mitigate the influence of the exponential increase in saturation values, we considered the pause sum at 90th percentile as maximum occupancy signal in datasets without UMIs. Using the maximum occupancy value, we scaled all pause and gene body densities from 0 to 1 to estimate initiation rates (Eq. 12). The assumption that the pause sum saturates at the 90th percentile of ranked pause sums in datasets without UMIs may not be accurate, as the pause sum at the 90th percentile is lower than the pause signals observed for heat shock activated genes (such as *Hsp70*) in *Drosophila* (Fig. S3B&C). Although our initiation rate calculations are based on these assumptions, more accurate estimates can be obtained by calibrating saturation values using datasets with known RNA polymerase densities in specific genomic regions.

## Data Availability

A Python package is currently installable from PyPi: https://pypi.org/project/compartmentModel using the command: `python3 -m pip install compartmentModel`.

The details of installation, processing of sequencing data, running the model, generating the figures, and UCSC Genome Browser track hubs are available this vignette: https://guertinlab.github.io/compartment_model/compModel_vignette/compartmentModel_vignette.html.

## Results

### Direct inhibition of pause release and initiation verifies the model predictions

The compartment model we developed interprets the mechanistic steps of RNA polymerase initiation, pause release, premature termination, and elongation as rate constants that explain observed changes in nascent transcription after acute perturbation and treatments. To validate the model predictions for changes in pause release rates (Eq. 8), we analyzed GRO-seq data following the inhibition of pause release with flavopiridol (11). Flavopiridol is a kinase inhibitor that targets CDK9 (65, 66), a key component of the positive transcription elongation factor b (P-TEFb) complex. P-TEFb promotes the release of paused RNA polymerase II into productive elongation, partially through the phosphorylation of the RPB1 C-terminal domain (4, 65, 67). Treatment with flavopiridol (0.3 µM) for 25 minutes results in a decrease of the pause release rate for ~82% of repressed genes, with a median rate decrease of ~57% (Fig. 2A, Fig. S4, & Fig. S5A). Additionally, we observed a modest reduction in the initiation rate, depending on the relative rates of premature termination and pause release (Fig. 2B).

**Fig. 2.**
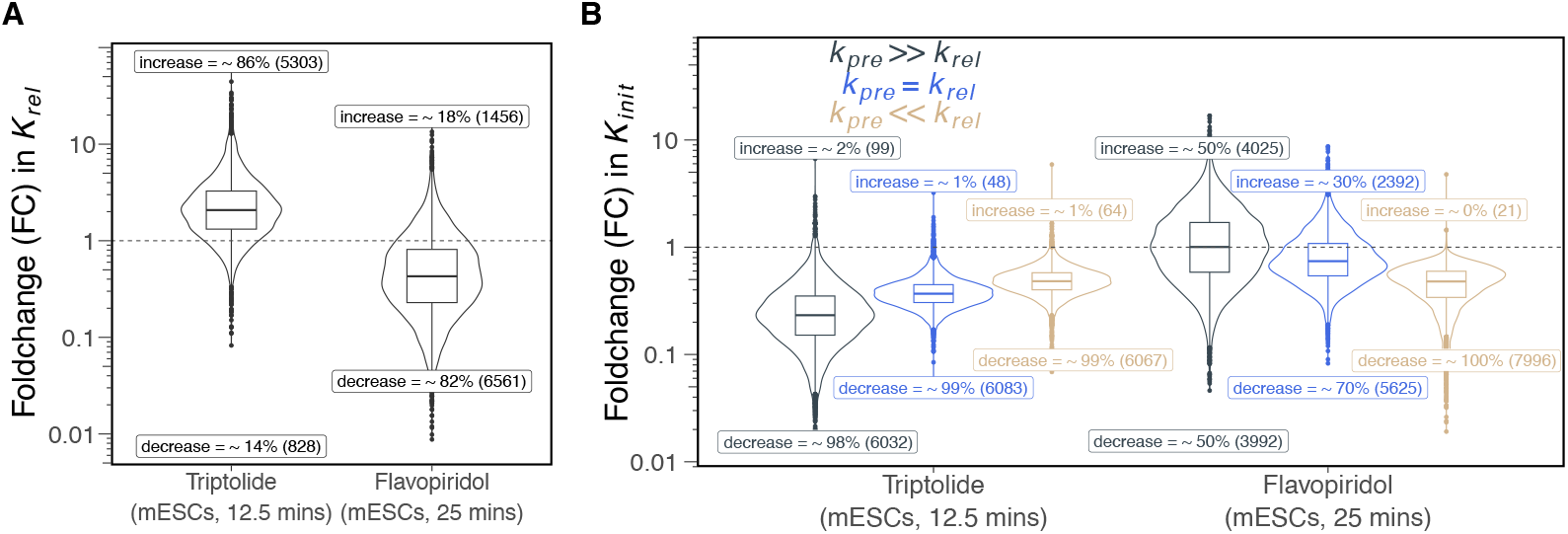
Inhibition of initiation and pause release validate the model. A) Flavopiridol treatment decreases pause release for ~82% of repressed genes, consistent with its known inhibitory effect on CDK9. Inspection of genes with increased pause release reveal a decrease in pause density for all genes (Fig. S4A-C). This unexpected reduction in pause density is attributed to spiky signals in the pause region or insufficient reads to accurately define the pause window (Fig. S4D-I). Triptolide increases pause release at most genes. Although this was not expected, this phenomenon could be a result of a general compensatory mechanism to increase transcription when initiation is inhibited. B) The bounds on changes in initiation rate are estimated with distinct relative values of pause release (*k*_*rel*_) and premature termination rate (*k*_*pre*_) (Materials & Methods). As expected, nearly all repressed genes (>99%) decrease their initiation rate upon triptolide treatment. The median fold change in *K*_*init*_ for TRP-repressed genes were 0.23 (*k*_*pre*_ >> *k*_*rel*_), 0.37 (*k*_*pre*_ = *k*_*rel*_), and 0.48 (*k*_*pre*_ << *k*_*rel*_). Flavopiridol does not show any clear effect on initiation if *k*_*pre*_ >> *k*_*rel*_, but appears inhibitory if *k*_*pre*_ << *k*_*rel*_ or *k*_*pre*_ = *k*_*rel*_.

The model links changes in a gene’s initiation rate to changes in body and pause densities. The change in initiation rate for a gene is constrained by the fold change in either pause density or body density, depending on the relative rates of premature termination and pause release (Eqs. 11 & 12). To validate the model predictions, we analyzed genome-wide nascent transcriptome data after acute treatments with an inhibitor of initiation, triptolide (TRP)(11). Triptolide blocks transcription initiation by inhibiting the ATPase activity of XPB (69), an ATP-dependent DNA translocase and a core component of TFIIH (70, 71). Treatment with 0.5 µM triptolide for 12.5 mins (11) causes the expected decrease in initiation rate for more than 98% of repressed genes, with a median decrease of ~77% when premature termination is faster than pause release (*k*_*pre*_ >> *k*_*rel*_, Fig. 2B, Fig. S5B). When pause release is comparable to or faster than premature termination (*k*_*rel*_ = *k*_*pre*_ or *k*_*rel*_ >> *k*_*pre*_), triptolide treatment leads to a median decrease in initiation rate of approximately 63% or 52% (Fig. 2B). Since initiation rate changes are constrained within a range and are dependent on relative rates of premature termination and pause release, we examined the results more closely to better understand these relative rates.

### Inhibition by triptolide indicates that premature termination is faster than pause release

Since changes in initiation rate are defined by relative values of premature termination and pause release (Eqs. 11–12), we sought to determine which rate is likely to be faster. To quantify the relationship between *k*_*pre*_ and *k*_*rel*_, we considered changes in the initiation rate (Eq. 6) for repressed genes after triptolide treatment (Fig. 2B). Specifically, we reasoned that the fold change in initiation rate for all repressed genes will be close to 0.25, based on the results that initiation is decreased by ~75% after 10 minutes of 1µM triptolide treatment (55). For each gene that has fold change in initiation rate range that spans 0.25 (Fig. S6), we calculated the ratios of premature termination rate to control pause release rate (Fig. 3A, Eq. 17). The median 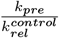 ratio is 2.6, with an inter-decile range of 0.8 - 13.4 (Fig. 3A). Premature termination is faster than pause release for over 85% of the genes (Fig. 3A). This trend is consistent among genes that are classified according to their expression level (Fig. S7). This suggests that the rate of premature termination is much faster than pause release at the majority of gene promoters, and on average only one out of every four paused polymerases proceeds into productive elongation. These results are consistent with previous data that suggests premature termination rates are faster than pause release rates (9, 55, 72–75) (Table S2).

**Fig. 3.**
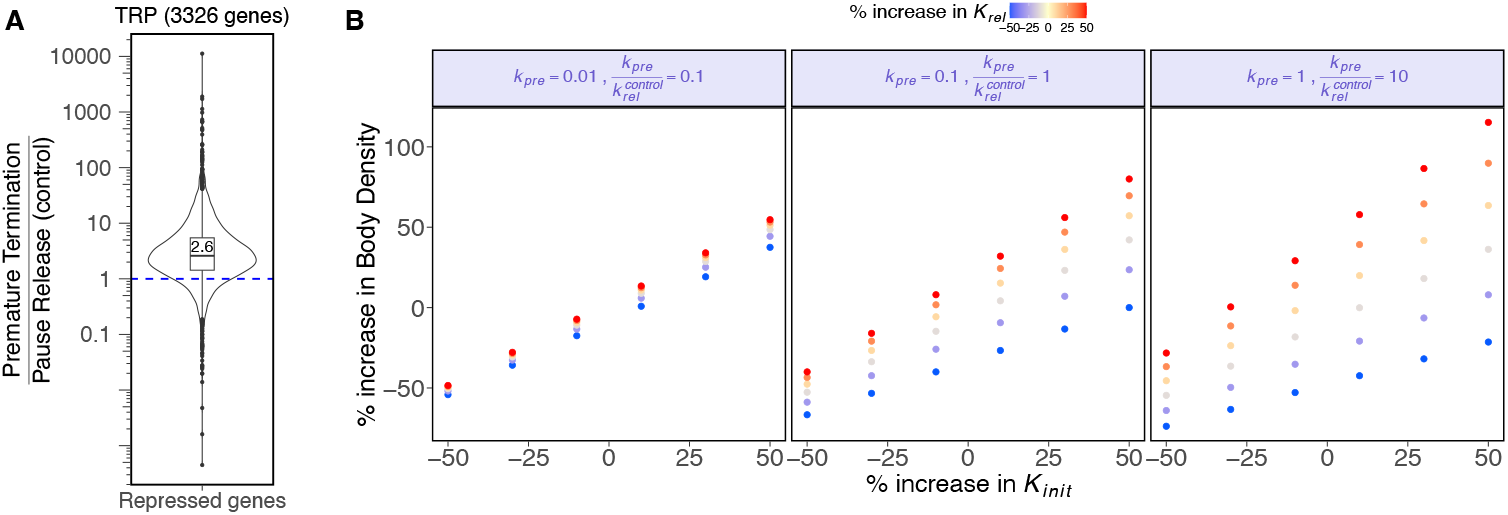
Experimental data and the model equations indicate that premature termination is faster than pause release. A) We set all TRP-repressed genes to have fold change (FC) in initiation rate of 0.25 if the gene’s limits of FC(*k*_*init*_) spanned this value (Eq. 11–12) (Fig. S6), then calculated each gene’s ratio of premature termination rate and pause release rate (Eq. 17). B) Changes in pause release affect changes in body density (transcription output) when premature termination *k*_*pre*_ is faster (*k*_*pre*_ >> *k*_*rel*_). When *k*_*pre*_ is relatively slower than *k*_*rel*_, change in body density is closely dependent upon changes in *k*_*init*_ and is minimally affected by changes in *k*_*rel*_. As termination *k*_*pre*_ becomes faster than pause release *k*_*rel*_, the body density becomes dependent on changes in *k*_*rel*_ and approaches the product of changes in initiation *k*_*init*_ and pause release *k*_*rel*_ rates. Pausing is regulated to control expression output in metazoans (68), so these results support premature termination being faster than pause release at most genes.

The result of flavopiridol inhibition on pause release also suggests that *k*_*pre*_ >> *k*_*rel*_. The pause inhibition data was generated with 300 nM flavopiridol (FP) for 25 minutes, which should cause a decrease in pause release without preferentially stimulating or inhibiting initiation. However, our analysis indicates that if *k*_*pre*_ << *k*_*rel*_, then nearly every gene would have to decrease initiation after FP treatment to explain the observed changes in pause and gene body densities (Fig. 2B) (11). While flavopiridol may interact with other kinases, such as CDK7 (66), at this concentration flavopiridol is unlikely to cause a decrease in initiation at nearly all genes. Moreover, the observed increase in RNA polymerase density at pause sites following flavopiridol treatment (11) contrasts with the decrease in pause density reported for CDK7 inhibition (76). Both the triptolide and flavopiridol analyses support the conclusion that premature termination occurs at a rate substantially faster than pause release.

### Change in pause release affects transcription output when premature termination is faster than pause release

To better understand how the rates of initiation, pause release, and premature termination collectively influence gene expression, we examined how changes in these rates determine transcription output. We use RNA polymerase density in the gene body as a proxy for change in transcription output. The change in gene body density is determined by the product of three quantities: (a) the fold change in initiation rate (*k*_*init*_), (b) the fold change in pause release rate (*k*_*rel*_) and (c) the inverse of fold change in “net exit rate” from the pause region, 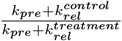. Depending on the relative values of pause release (*k*_*rel*_) and termination rate (*k*_*pre*_), the inverse of fold change in exit rate 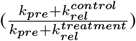 ranges from 1 (*k*_*pre*_ >> *k*_*rel*_) to the inverse of fold change in pause release (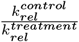 when *k*_*pre*_ << *k*_*rel*_) (Eqs. 9–10).

Thus, when pause release is much faster compared to termination (*k*_*rel*_ >> *k*_*pre*_), the change in transcription output closely mirrors the change in initiation rate and variations in pause release (*k*_*rel*_) have minimal impact on transcription output (Fig. 3B left panel). As *k*_*pre*_ becomes faster than *k*_*rel*_, changes in pause release (*k*_*rel*_) become more influential, and transcription output approaches the product of change in initiation and change in pause release rate (Fig. 3B right panel). These results demonstrate that regulating pause release has a bigger effect on gene expression output when premature termination occurs at a relatively faster rate.

These analyses, taken together with the analysis of pause-inhibition and initiation-inhibition data, as well as previous work (Table S2), suggests that premature termination is faster than pause release at most genes. Therefore, we maintain this assumption throughout the main figures and text, meaning that changes in initiation rate are interpreted as fold change in pause densities as measured by PRO-seq (Eq. 11).

### Mechanistic functions of transcription factors

To determine the regulatory roles of transcription factors, we analyzed existing PRO-seq datasets that rapidly degraded TBP and ZNF143 (43, 44). As expected from TBP’s established role in initiation (77), degradation of TBP reduces the initiation rates for more than ~86% of repressed genes (Fig. 4A, Fig. S8A, Fig. S9). Previous work proposed the role of ZNF143 as an initiation factor by demonstrating that ZNF143 influences transcription start site (TSS) selection, with alternative TSSs being utilized in repressed genes following ZNF143 degradation (44). We confirm that the degradation of ZNF143 led to a decrease in initiation rates for ~87% of repressed genes (Fig. 4A, Fig. S8B, Fig. S9).

**Fig. 4.**
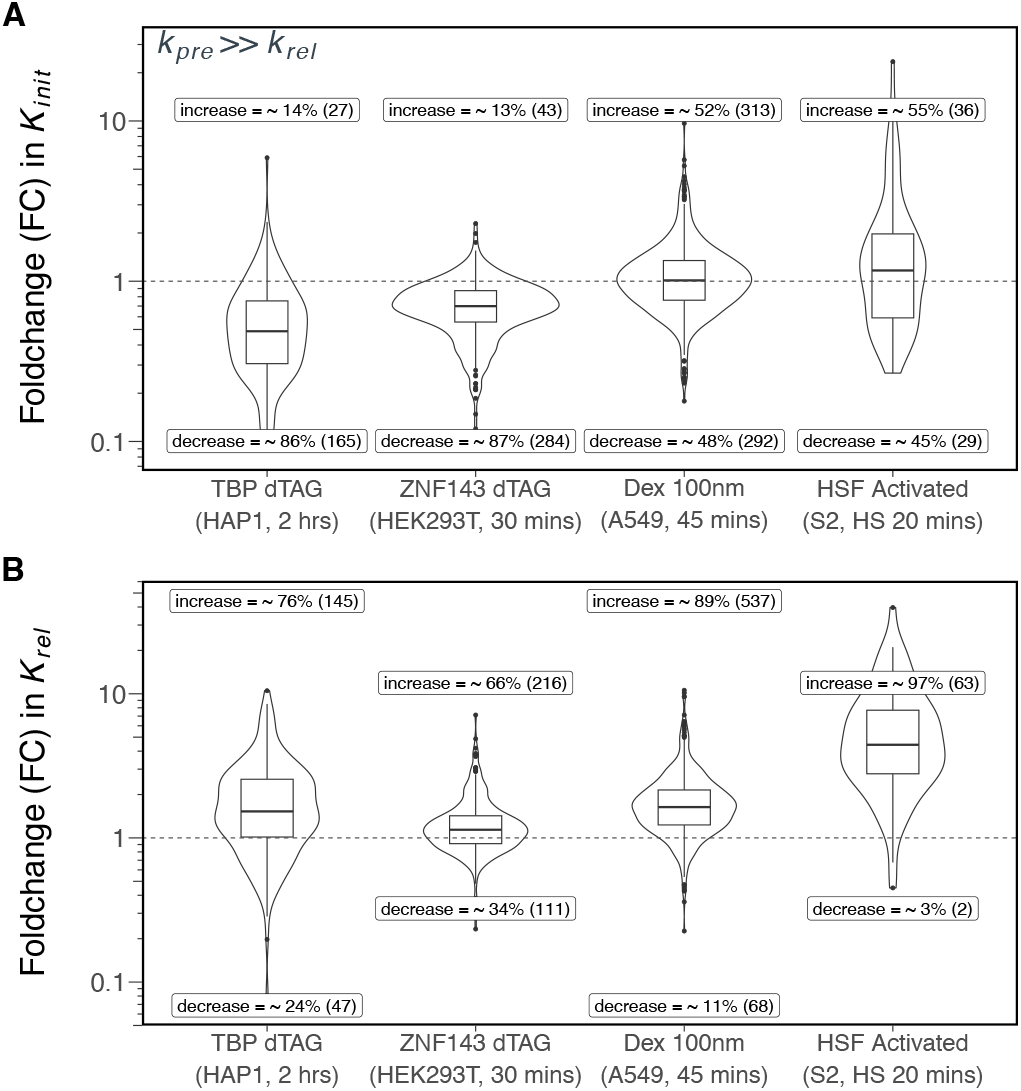
Modeling kinetic PRO-seq data determines the regulatory roles of activated and degraded transcription factors. A) Degradation of ZNF143 (44) and TBP (43) reduces initiation for approximately 90% of repressed genes in each dataset. Dexamethasone-activated genes (45) and Heat Shock Factor (HSF)-activated genes in S2 cells (33) do not show a clear direction of change in initiation rate, suggesting that *Drosophila* HSF and human glucocorticoid receptor (GR) may not regulate initiation. Fig. S9 contains the estimated fold change in initiation rates if premature termination is less than pause release. B) Dexamethasone (45) enhances pause release in approximately 89% of activated genes, indicating that GR’s role is to regulate pause release. In *Drosophila* S2 cells, HSF activates target genes by facilitating pause release. Degradation of TBP and ZNF143 tends to increase pause release for repressed genes, similar to the effect seen in repressed genes after triptolide treatment (Fig. 2.)

Next, we investigated the function of glucocorticoid receptor (GR) by analyzing PRO-seq data after acute dexamethasone (Dex) treatment (45). Upon binding dexamethasone, GR translocates to the nucleus, binds DNA and directly activates transcription (78–80). After 45 mins of dexamethasone treatment in A549 cells (45), pause release rate increased for 89% of activated genes (Fig. 4B, Fig. S8C) with no clear change in initiation rate for Dex-activated genes (Fig. 4A, Fig. S9), indicating that GR predominantly activates its target genes by stimulating productive release of paused RNA polymerases. We further confirmed that dexamethasone activation of GR stimulates pause release in U2OS and C7 cells (45, 46, 81) (Fig. S10).

Promoter-proximal pausing was first characterized at *Drosophila* heat shock genes (2) and the *Drosophila* heat shock factor (HSF) is known to stimulate pause release (33, 82, 83). Our analyses confirms that nearly all (~97%) HSF-activated genes in *Drosophila* S2 cells show an increase in pause release upon heat shock (Fig. 4B, Fig. S8D). Taken together, these analyses of existing kinetic PRO-seq data demonstrates the utility of this approach when assigning molecular functions to transcription factors.

### Inferring absolute and effective pause release rates

Our model estimates the absolute values of pause release rates as the product of elongation rates and the inverse of a gene’s *pausing index* (Eq. 13). Since these estimates are derived from the ratio of body to pause densities, they are independent of the scaling or normalization criteria applied to genomic data. We considered 2000 bp/min a representative elongation rate and we kept this constant for all genes because we cannot estimate accurate gene-specific elongation rates. We report the pause release rates before and after dexamethasone treatment because GR regulates pause release (Fig. 4B & Fig. S10). The median pause release rate constants increase from 0.86 events/min to 1.35 events/min (Fig. 5A).

**Fig. 5.**
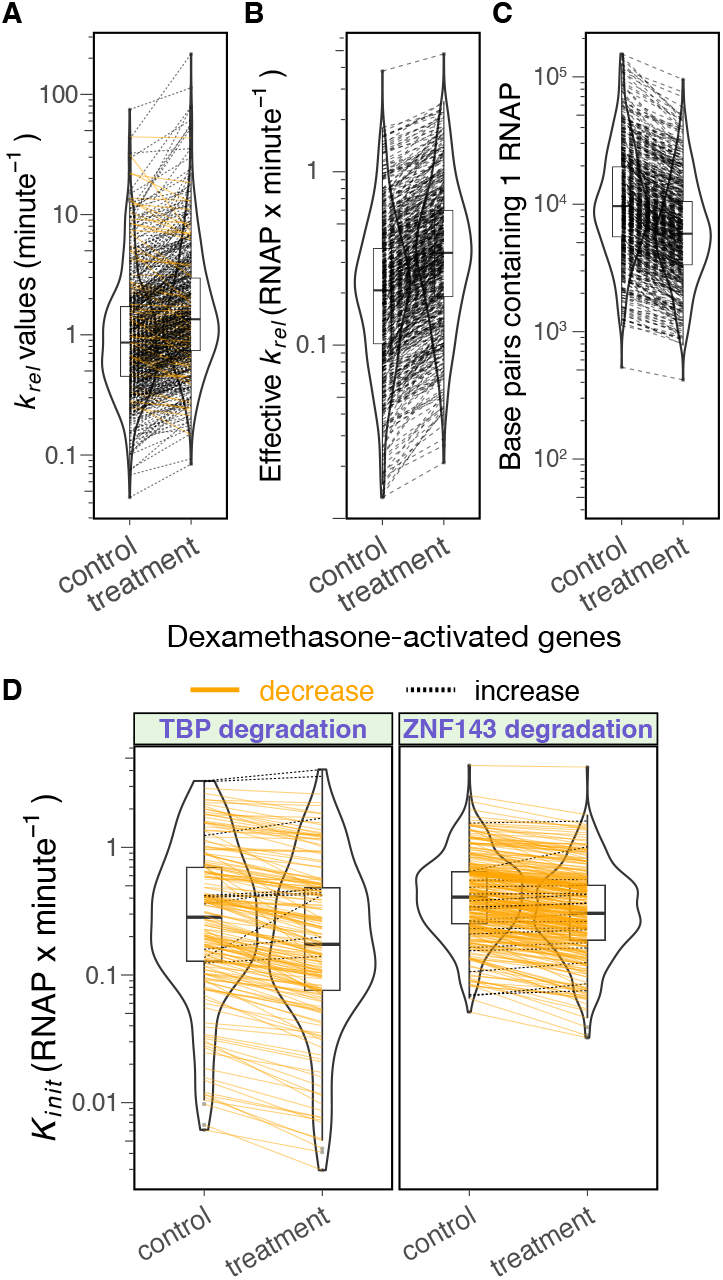
Rates of initiation and pause release can be estimated using steady-state densities of pause and gene body regions. A) The median pause release rates of Dex-activated genes increase from 0.86 events/min (control) to 1.35 events/min, with an inter-decile range of 0.27 - 4.1 events/min in untreated conditions and 0.42 - 7 events/min after dexamethasone treatment. 89% of the Dex-activated genes (537 out of 605) increase their pause release rate. B) The effective pause release rate represents the release of RNA polymerase into the gene body and is calculated as the product of the pause release rate constant and the scaled pause density (Fig. S2E). The median effective pause release rate for Dex-activated genes increases 1.6-fold, from 0.21 RNAP/min to 0.34 RNAP/min with an inter-decile range of 0.06 - 0.63 RNAP/min under untreated conditions and 0.09 - 1 RNAP/min after dexamethasone treatment. C) The density of RNA polymerase in gene bodies increased after dexamethasone treatment with median density increasing from 1 RNAP every 9.6kb to 1 RNAP every 5.8kb. The inter-decile ranges were 1 RNAP every 3.1 kb - 36.3 kb in untreated conditions, increasing to 1 RNAP every 1.8 kb - 20.3 kb after dexamethasone treatment. D) Degradation of TBP in HAP1 cells decreases the median initiation rates from 0.3 RNAP/min (inter-decile range of 0.04 - 1.6) to 0.17 RNAP/min (inter-decile range of 0.03 - 1.2 RNAP/min). Degradation of ZNF143 in HEK293T cells decreases the median initiation rates from 0.41 RNAP/min (inter-decile range of 0.16 - 1.1 RNAP/min) to 0.3 RNAP/min (interdecile range of 0.11 - 0.8 RNAP/min).

We also estimated an effective pause release rate, *k*_*rel*_ × *P*, representing the number of polymerases entering productive elongation at steady state. Since calculating effective pause release requires an estimation of pause densities as absolute occupancy values, we used a saturation fit to scale pause densities (Fig. S2E) (Materials & Methods). Upon dexamethasone treatment, the median effective pause release rate of Dex-activated genes increased 1.6-fold from 0.21 RNA polymerase (RNAP) /min to 0.34 RNAP/min (Fig. 5B). Effective pause release rates increase for all Dex-activated genes (Fig. 5B). Ninety percent of Dex-activated genes have ≤ 1 RNAP entering the gene body per minute (Fig. 5B).

Next, we calculated RNA polymerase occupancy levels in the gene body of Dex-activated genes in both conditions (Eq. 15). Based on the effective pause release rates, the median estimated occupancy in untreated cells was 1 RNA polymerase per 9.6 kilobases (kb) (Fig. 5C). The density of RNA polymerase in the gene body increased after dexamethasone treatment to median of 1 RNA polymerase every 5.8 kb (Fig. 5C). These results align with previous imaging studies showing that two-thirds of nascent RNA transcripts are spaced more than 12 kb apart from another RNA polymerase, while the remaining 33% of transcripts exhibit densities ranging from 1 RNA polymerase per kb to 1 RNA polymerase per 8 kb (61, 62). Previously reported “initiation” rates of 0.16–0.4 events/min in single-molecule imaging studies (63) can be interpreted as effective pause release rates, corresponding to a body density of 1 RNA polymerase every 5–12.5 kb. Another study estimated effective pause release rates of 0.19–0.29 events/min, yielding a body density of 1 RNA polymerase every 6.9–10.5 kb (64). Our calculated pause release rates and genomic estimates of RNA polymerase occupancy align with independent imaging and single-molecule studies, underscoring the validity of our approach in converting unitless genomic RNA polymerase profiling values to absolute RNA polymerase occupancy estimates.

### Inferring absolute initiation rates

The initiation rate estimate incorporates premature termination rates, elongation rates, with pause and body densities as occupancy values (Eq. 14) (Materials & Methods). We concluded faster premature termination rates (*k*_*pre*_) than pause release rates (*k*_*rel*_) at gene promoters using experimental data from triptolide-treated cells (Fig. 3A). For the rapid degradation of TBP and ZNF143 datasets, we assumed that premature termination is 2.6 times faster than pause release (Fig. 3A). Upon TBP degradation in HAP1 cells, the median initiation rate decreased from 0.3 events/min to 0.17 RNAP/min (Fig. 5D). While, upon ZNF143 degradation in HEK293T cells, the median initiation rate decreased from 0.4 RNAP/min to 0.3 RNAP/min (Fig. 5D). These calculations indicate that the distribution of initiation rates and effective pause release rates fall within similar ranges.

### Most paused polymerases turn over in less than a minute

The residency time of a paused RNA polymerase is determined by the rates of premature termination and pause release. The median half-lives of paused RNA polymerases, calculated as ln (2)*/*(*k*_*pre*_ + *k*_*rel*_), across all treatments was approximately 16.6 seconds (Fig. 6), with an inter-decile range of 3.3 to 76 seconds, indicating that most paused polymerases turn over within a minute. Premature termination rates primarily determine half-lives rather than pause release rates. For instance, at *k*_*pre*_ = 10 × *k*_*rel*_ (Fig. S11), the turnover time and interdecile range decrease by an approximate factor of 3.8 (i.e., 10*/*2.6).

**Fig. 6.**
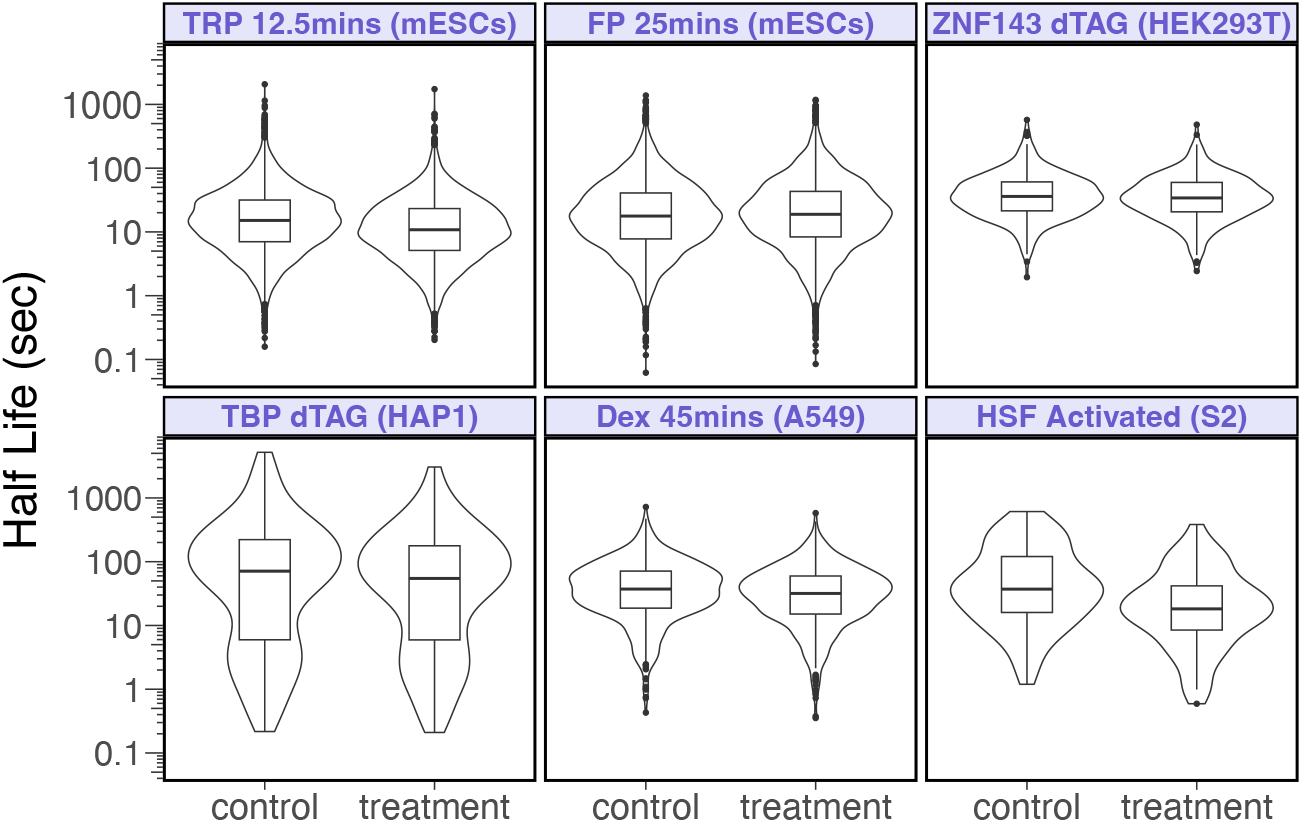
Most paused polymerases turn over rapidly. Assuming premature termination rates at each gene promoter as 2.6 times faster than pause release rates (Fig. 3A), the median half-life of paused polymerases across all datasets and treatments is 16.6 seconds, with an inter-decile range of 3.3 to 76 seconds. For repressed genes under triptolide treatment, the median half-life decreased from 15.1 seconds to 10.8 seconds. For HSF-activated genes, the median half-life decreased from 37.2 seconds to 18.2 seconds. For repressed genes after TBP degradation, the median half-life decreased from 71.1 seconds to 55 seconds. The half-lives in other datasets showed only minor changes after treatment.

Several studies are consistent with our estimates of short lived paused polymerases (9, 55, 73–75). A dual-enzyme single molecule footprint study performed triptolide and flavopiridol treatments to conclude that paused RNA Pol II half lives were <2.5minutes, which was the earliest time point measured (74). Another study photobleached RNA Polymerase II in live cells and modeled the results to conclude that the residency time of a paused RNA Polymerase II was approximately 42 seconds (9, 73). An independent study inhibited premature termination and pause release with hydrogen peroxide and flavopiridol, followed by nuclear walk-on assays, to estimate that a paused polymerase is replaced every 40 seconds (55). Another study used transcription inhibitors and ChIP to conclude that over 80% of paused RNA polymerases turn over in less than 2 minutes (75). Other studies have reported longer half-lives of paused polymerases over several minutes (10, 11, 72, 84). In the discussion we provide possible explanations for the disparate estimates in an attempt to reconcile seemingly inconsistent findings.

### Parameter sensitivity analysis supports the robustness of our conclusions

We based our rate estimates and conclusions on several assumptions that are grounded in experimental data: (1) premature termination and elongation rates are unaffected by triptolide treatment; (2) the average gene elongation rate is approximately 2kb/min; (3) 12.5 minutes of 1µM triptolide inhibits initiation by 75%; and (4) we can estimate the absolute density of RNA polymerases in the pause region. We varied parameters that represent these assumptions over a wide range to determine their influence on elongation rates, rate ratios, and our overall conclusions.

We investigated how varying premature termination and elongation rates affect the ratio of premature termination to pause release (Eq. 16, Fig. S12). If triptolide treatment decreases the elongation rate of elongating RNA polymerases by more than 20%, then the majority of genes do not exhibit a faster premature termination rate compared to pause release rate (Fig. S12). We have no reason to expect that triptolide reduces the elongation rate of RNA polymerases in the gene body.

In the previous sections we assumed that the average elongation rate is 2kb/min. However, elongation rates can vary between genes and within a gene (34, 50, 85, 86). Our analysis ignores the fact that elongation rate varies within a gene because we average the gene body density, which effectively averages the elongation rate in our model. We varied the average elongation rate from 1-5kb/min to determine how elongation speed influences other rates and our conclusions. In short, changing the elongation rate causes linear and proportional changes in all other rates (Fig. S13), since pause release is linearly dependent on elongation (Eq. 13) and premature termination is related to pause release (Eq. 17). For example, if we increase elongation rate by 2 fold, then all other rates will change by two-fold. Since we consider that elongation and premature termination rates remain unchanged after triptolide treatment, elongation rates do not influence *k*_*pre*_*/k*_*rel*_ (Eq. 17), fold changes in initiation (Eqs. 5 & 6), and fold changes in pause release (Eq. 6). However, faster elongation rates would shorten the residency times of paused RNA polymerase due to faster termination and pause release (Fig. S13).

We assumed that 12.5 minutes of 0.5 µM triptolide caused an average fold-change in the initiation rate of 0.25 for triptolide-repressed genes in mESC cells (11). This estimate is based on an *in vitro* estimate of genome-wide decrease in initiation after a time course (1, 3, 10 and 30 mins) of 1µM triptolide treatment (55). Uncertainty in this parameter will affect our estimates of absolute premature termination rates, absolute initiation rates, and the premature termination to pause release ratio (Fig. S14). If triptolide treatment is less effective than we assume (i.e. fold-change in initiation is >0.25), then the premature termination and initiation rates decrease (Fig. S14C&D, Eqs. 14 & 17). The pause release rates for gene sets at different efficacies of triptolide inhibition show modest differences (Fig. S14B). If triptolide causes a modest 0.35 fold-change in initiation, then approximately half of genes have a *k*_*pre*_ > *k*_*rel*_ (i.e. termination and pause release rates are comparable) (Fig. S14A). The reciprocal is also true: if we are underestimating triptolide efficacy, then premature termination of a paused RNA polymerase is more likely than we report. Our estimate of triptolide efficacy is based on experimental data and 75% inhibition spans the empirically determined bounds of initiation rate fold change (Fig. S6), but other studies assume that this same triptolide treatment is nearly complete and more rapid (11). Although we only focus on triptolide-repressed genes herein, all genes are likely differentially affected by triptolide treatment, some completely inhibited and others insensitive. In the future, gene-specific measurements or estimates are needed to quantify the relationship between termination and pause release at individual genes.

We used a saturation function to estimate “full” occupancy of the pause region, which allowed us to calculate the absolute density of RNA polymerase in the gene body (Fig. S2A-F). Since the triptolide dataset (11) doesn’t have UMIs, we had considered 90th percentile as saturation to be consistent with saturation values observed in datasets with UMIs (Fig. S2D-F). To investigate how saturation values may affect occupancy, we varied them by 5-fold in each direction (Fig. S15). Based on pause signal of *x* arbitrary units at the 90th percentile, we considered pause sums at 48, 70, 90, 95, 99.8 percentiles of ranked pause sums (P = 0.2*x*, 0.4*x, x*, 1.8*x* and 5*x*). The ratio of premature termination rate to pause release rate remains stable over this wide range of absolute RNAP density estimates (Fig. S15A), with modest <2.4-fold differences in both rates over the 25-fold range (0.2*x* to 5*x*) (Fig. S15A,B&C). Initiation rate is more sensitive to the RNAP density, decreasing proportionally as RNA polymerase density estimates increase (Fig. S15D).

Scaling absolute pause occupancy proportionally affects gene body density calculations. By inferring the RNAP density in the pause region, we can calculate the median values of RNA Polymerase density within genes. These median gene body densities equate to 1 RNAP every 1.4 kb (48 percentile; 0.2*x*), 1 RNAP every 5.3 kb (90 percentile; *x*), and 1 RNAP every 25.2 kb (99.8 percentile; 5*x*) (Fig. S15E). The 90th percentile saturation values (Fig. 5C, Fig. S15E) equate to gene body densities that are supported by multiple imaging studies: 1 RNAP every 8-12 kb (61), 1 RNAP every 5-12.5 kb (63) and 1 RNAP every 6.9-10.5 kb (64). Although, absolute RNAP occupancy estimates influence all calculations except the *k*_*pre*_*/k*_*rel*_ ratio, defining “full” occupancy using the 90th percentile of pause density, guided by saturation curve estimates (Fig. S2), yields gene body densities that align best with independent RNA polymerase occupancy measurements.

## Discussion

The results of this study provide insights into the kinetic mechanisms of transcriptional regulation, with a focus on the fate of paused RNA polymerases. Our modeling framework, validated with kinetic PRO-seq datasets, reveals that most paused polymerases terminate prematurely rather than progressing into productive elongation (Fig. 7 and Table S3). We found that the rate of premature termination (*k*_*pre*_) is significantly faster than the rate of pause release (*k*_*rel*_) for most genes. By leveraging triptolide inhibition data, we estimated the ratio of premature termination to pause release rates (*k*_*pre*_*/k*_*rel*_) and found that premature termination occurs approximately two and half times more frequently than productive elongation. The relative rates of termination versus pause release has significant implications for understanding transcriptional control. Transcription factors that enhance pause release, such as HSF and GR, have greater influence on gene expression in contexts with fast termination rates. The ability of our model to quantify these rates provides a framework for determining how different factors mechanistically regulate transcription.

**Fig. 7.**
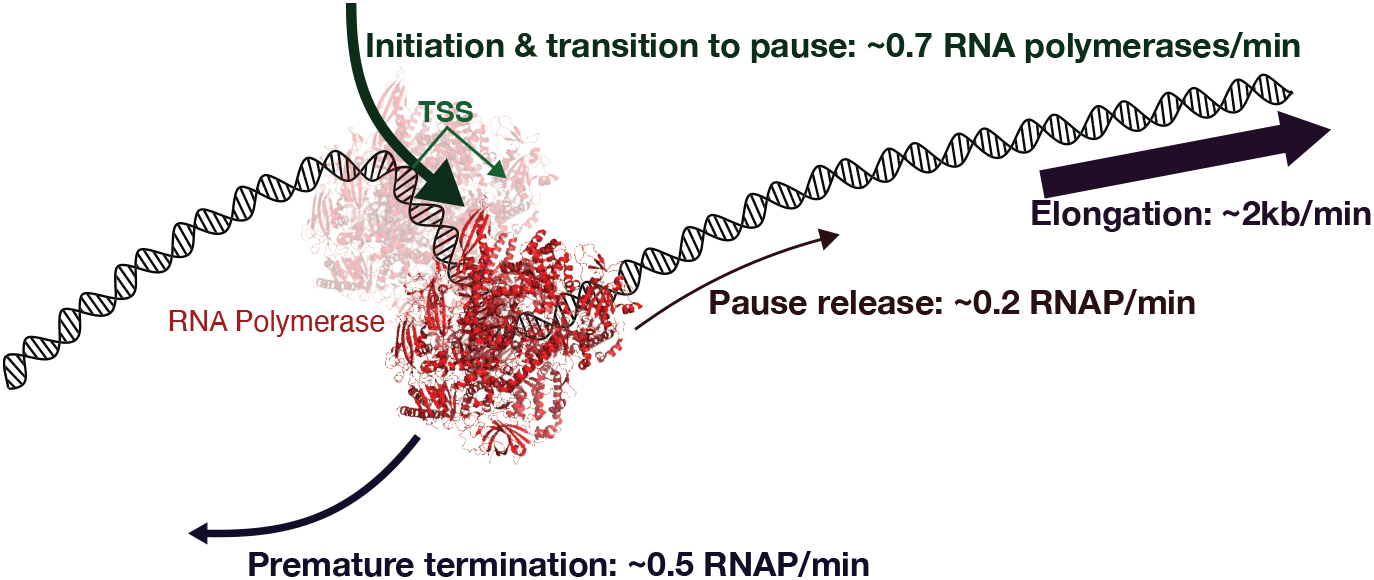
A kinetic model of initiation, termination, and early elongation. The cartoon represents the conclusions from our modeling of kinetic nascent transcriptome data. The initiation rate of 0.7 RNA polymerases (RNAP) per minute assumes a non-limiting amount of RNA polymerase available to initiate. The translucent RNA polymerase represents pre-initiation or abortive initiation complexes; so the term *initiation* rate within our model refers to the series of events that have to occur for an RNA polymerase to successfully proceed into the pause region. The average effective pause release rate is 0.2 RNAP/minute and the average effective termination rate of pause RNA polymerases is approximate 2.6 times faster at 0.5 RNAP/minute. These rates are derived from well established rates of elongation (~2 kilobases/minute) averaged over a gene body, experimental triptolide efficacy data, and empirical estimates of fully occupied pause sites. The effective rates in different datasets are consistent with each other (Table S3).

Our framework was validated using datasets in which initiation and pause release were directly inhibited. The effects of triptolide on initiation and flavopiridol on pause release were consistent with known molecular mechanisms, demonstrating that our model accurately captures kinetic changes in transcription. Furthermore, the model successfully classified transcription factors based on their regulatory roles, confirming TBP and ZNF143 as initiation factors and HSF and GR as regulators of pause release.

### Model limitations

Sensitivity analyses further support the robustness of our conclusions. While absolute values of rates depend on estimates of elongation speed and triptolide efficacy, the key finding that termination is faster than pause release remains consistent across a broad range of estimates. Moreover, our calculation of gene body RNA polymerase densities from the modeling align with previous imaging studies, reinforcing the validity of our RNA polymerase occupancy estimates.

One limitation of our current framework is that it does not explicitly take transcriptional bursting into account. Transcription occurs in stochastic bursts, where genes switch between active and inactive states, leading to fluctuations in RNA polymerase occupancy (87). Although our model assumes steady-state transcription rates, it can be adapted to incorporate bursting kinetics if the burst frequency and duration are known for a given gene. For example, if 20% of a gene’s alleles are active, then the saturation curve would be re-calibrated so that complete occupancy would be considered 20% of the estimated full occupancy. When we consider bursting and the fact that only a fraction of the genes in a population are active, changing saturation values affects initiation rates (Eq. 14) by approximately the same scaling factor (Fig. S15D). The half-lives and pause residency time at genes are independent of bursting activity because pause release and premature termination rates do not depend on absolute RNA polymerase levels (Eqs. 13, 16 & 17). Future work could integrate live cell imaging or single molecule footprinting data to refine the model and more fully capture dynamic transcriptional regulation for each gene.

### Reconciling disparate estimates of rates and pause residency times from previous studies

We and others have found that premature termination is much faster than pause release (Fig. 3A) and paused RNA polymerases turn over rapidly (Fig. 6) (9, 55, 73–75).

Some previous studies report slower pause release and premature termination rate constants (Table S2) (72), with estimated half-lives in the range of several minutes (72, 84). A key distinction between these studies and our approach is that they exclusively measured capped short nascent RNA (72, 84), while we quantify all engaged RNA polymerases that are competent to elongate (88, 89). We speculate that paused RNA polymerases with capped RNA are less likely to prematurely terminate than polymerases with uncapped RNA, potentially accounting for the observed differences. If nearly all initiated polymerases terminate prematurely (9) (Fig. S16), the slow turnover rates observed in (72, 84) may primarily reflect the termination dynamics of the minority population of paused RNA polymerases with capped RNA, with minimal contribution of pause release to measurements in both untreated and flavopiridol conditions.

Another study that reported longer half-lives in minutes had used 0.5µM triptolide treatment over several time points (11). Since the initial publication of this study, comparable times and concentrations of triptolide were shown to be insufficient to fully inhibit initiation (55). Our analysis also provides an upper bound for initiation inhibition, which supports these *in vitro* measurements of triptolide efficacy (55). Anything less than immediate and complete inhibition of initiation would cause an inflation of pause residency time because all newly initiated RNA polymerases will be considered stably paused.

A live cell imaging study used a photoactivatable GFP-RNA Pol II to measure pause stability at an uninduced and heat shock-induced *Hsp70* transgene (10). This work was performed in polytene chromosomes, averaging the RNAPII signal over a thousand copies of *Hsp70*. This analysis assumes that paGFP-RNAPII will dissociate from the locus upon termination, but if terminated RNAPII reinitiates at a proximal *Hsp70* gene, it will not contribute to the decay of GFP signal from the locus. This will result in an inflated half-life of paused polymerases because any local reinitiation will be considered stably paused. They also used a biochemical fractionation experiment upon inhibition of initiation with 10µM triptolide to quantify decay of nascent paused RNA (10). Again, subsequent studies found that even 10µM triptolide is not sufficient to immediately and completely inhibit initiation (55). This leads to a deflated decay rate because newly initiated RNA polymerases are indistinguishable from polymerases that are stably paused.

### Future Directions

Although our model is based on known mechanisms of initiation and early elongation and provides valuable insights into transcription dynamics, future work could refine these estimates using direct measurements of elongation rates at individual genes. Expanding this model to incorporate gene-specific variations in elongation and bursting kinetics would enhance its predictive power. Our current estimates of initiation rate encapsulate several molecular processes such as chromatin opening, pre-initiation complex assembly, abortive initiation, and promoter escape. As approaches become more sophisticated and can differentiate between these early stages, future models may be able to provide further mechanistic insights informing on these early processes.

By quantifying initiation, pause release, and termination rates, our framework and corresponding `compartmentModel` software are powerful tools for dissecting the kinetic mechanisms of transcription and determining the molecular functions of transcription factors.

## ACKNOWLEDGEMENTS

This work was funded by R35-GM128635 to MJG. We thank Pedro Mendes for modeling discussions and providing parameter estimations techniques.

## AUTHOR CONTRIBUTIONS

RM and MJG conceptualized and developed the project. RM and MJG developed the model. RM analyzed the data. RM developed the software. RM and MJG wrote the manuscript.

---

**Fig. S1.**
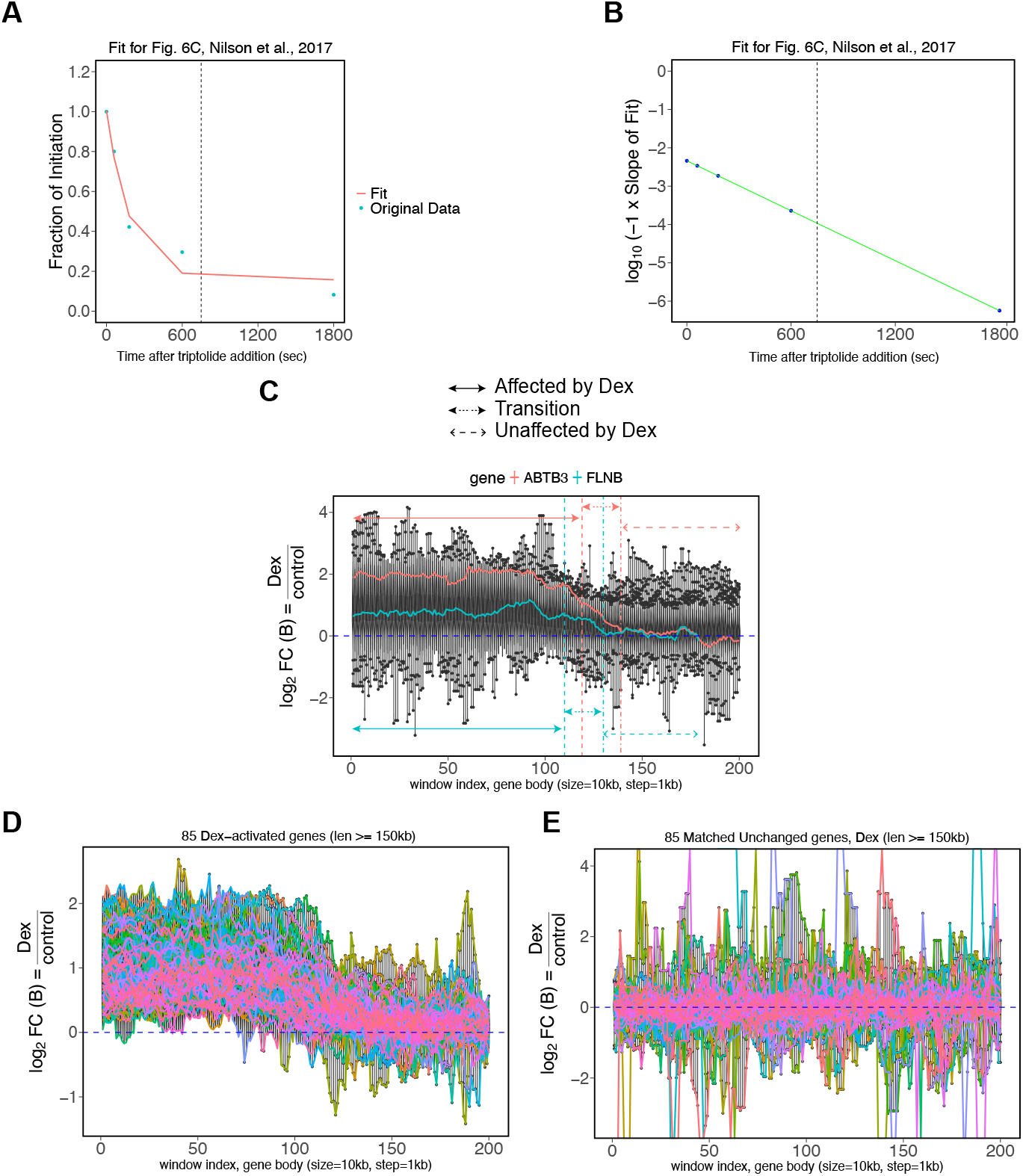
Validation of steady-state assumptions for triptolide and dexamethasone treatments. A) We fit an exponential decay curve (*a* ∗ *e*^−*b*∗*x*^ + *c*) to the time-series data for initiation following triptolide treatment (55). The dashed vertical line indicates T=12.5 minutes, the time point used in the current work. B) The slope of the exponential fit at T=12.5 minutes is approximately −10^−4^, validating the assumption of a steady-state initiation rate. C) Trend lines are shown for two representative genes with length ≥150kb. We observe a transition state between pre and post-Dexamethasone steady states. D) Eighty-five activated genes longer than 150kb have similar RNA polymerase distributions with two steady states and an intermediate transition state. E) Eighty-five unchanged genes (matched in length and expression to the activated genes in panel D) exhibit steady state profiles over the affected and unaffected regions.

**Fig. S2.**
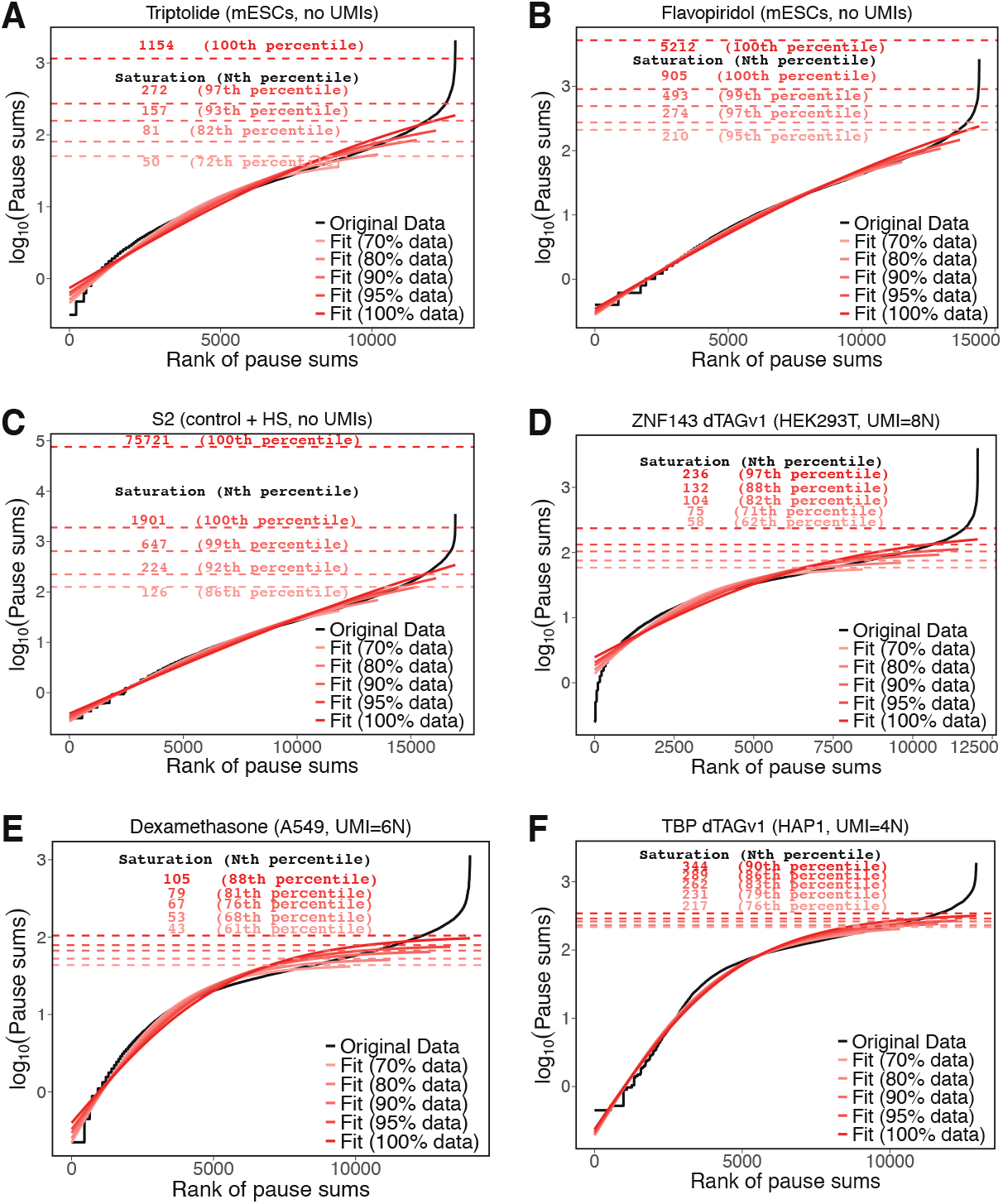
Saturation curves for datasets with Unique Molecular Identifiers (UMIs) provide estimates of RNA polymerase occupancy. The plots show the saturation curves and values from fits to the hyperbolic tangent function for datasets lacking Unique Molecular Identifiers (UMIs) (A-C) or containing UMIs (D-F). Datasets lacking UMIs do not have stable saturation values when ranges of pause sum percentiles are used to fit the saturation curve.

**Fig. S3.**
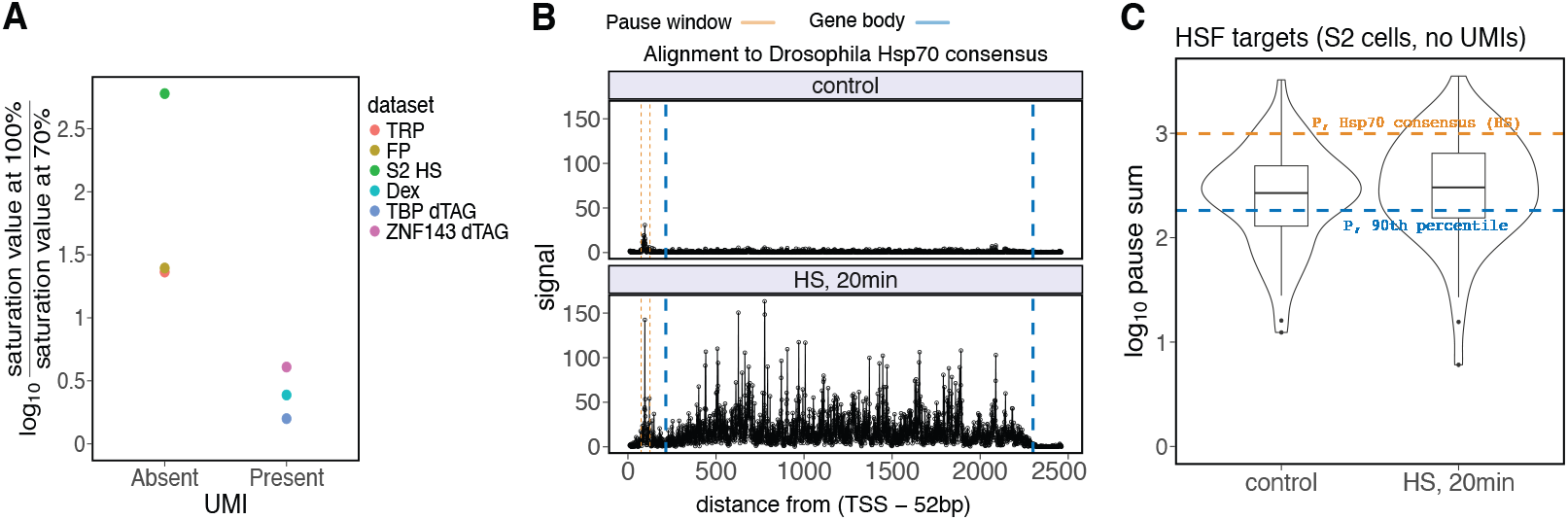
Saturation values from datasets lacking Unique Molecular Identifiers (UMIs) are unreliable as maximum occupancy signal of RNA polymerase. A) The ratio of saturation values derived from fits to 100% of the data compared to 70% of the data (ignoring the top 30% of pause sum genes) is substantially higher for datasets without UMIs, ranging from 23-fold to 600-fold. In contrast, datasets with UMIs show much lower variability, with ratios ranging from 1.6-fold to 4-fold. B) These plots show the normalized read signal for the *Hsp70* consensus sequence from the dataset (33) used in this study. C) The pause sum of *Hsp70* (P = 992) is five times higher than the pause sum of the gene at the 90th percentile (P = 182). The violin plot represents the pause sums for all HSF-activated genes, most of which are higher than the 90th percentile value. Considering that *Hsp70* has a fully occupied pause region during heat shock (90), the assumption that the 90th percentile pause sum represents the maximum occupancy signal is not supported in this dataset lacking UMIs.

**Fig. S4.**
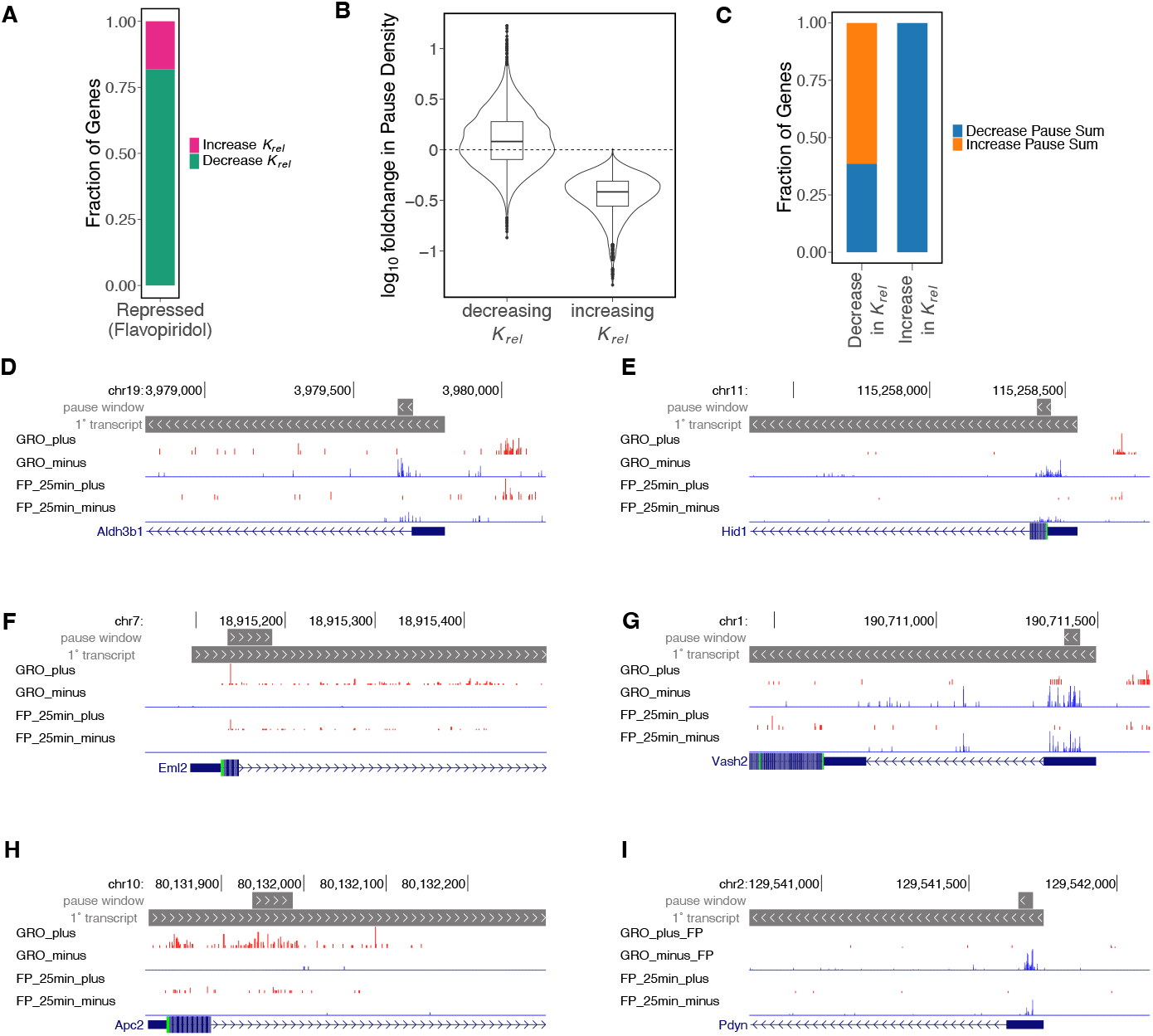
Genes with increased pause release after flavopiridol treatment have atypical read distributions. A) Approximately 82% of flavopiridol-repressed genes decrease their pause release rate (*k*_*rel*_). B) All genes with increased pause release rate also have a decrease in pause density. C) Sixty-one percent of repressed genes with decreased pause release have an expected increase in pause density after flavopiridol treatment. D-I) The UCSC Genome Browser snapshots show representative genes with increased pause release (*k*_*rel*_) and decreased pause density. High pause signals under untreated conditions in these genes may result from the lack of UMIs in the libraries (11).

**Fig. S5.**
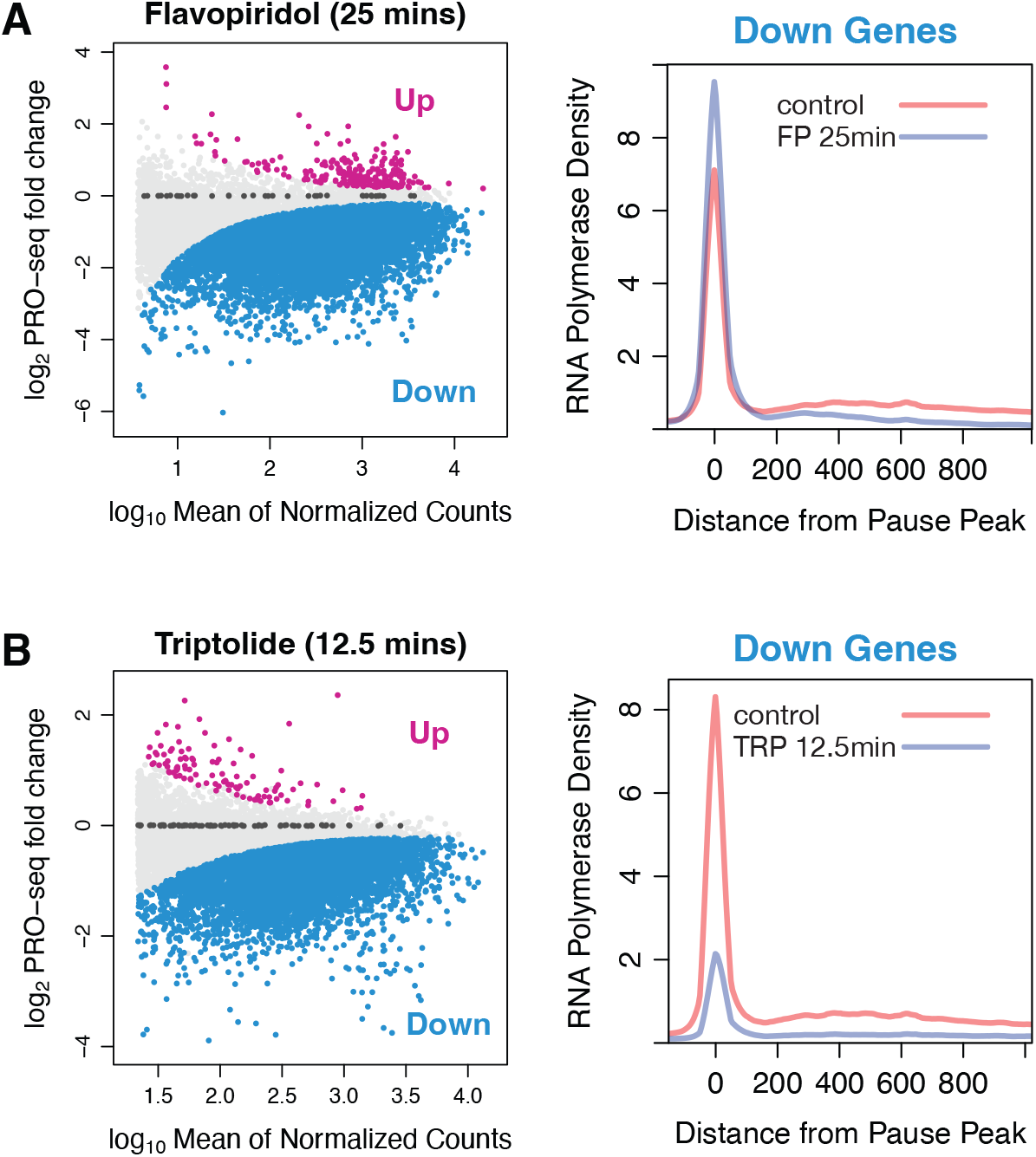
Flavopiridol and Triptolide induce genome-wide repression and changes in RNA polymerase distribution. A) Flavopiridol (FP) treatment for 25 mins (11) resulted in 12994 *Down* and 310 *Up* genes. Composite profiles of the *Down* genes show an increase in RNA polymerase density in pause window upon FP treatment. B) Triptolide treatment for 12.5 mins (11) resulted in 9632 *Down* and 104 *Up* genes. The *Down* genes have a decrease in RNA polymerase density in pause window and gene body upon triptolide treatment.

**Fig. S6.**
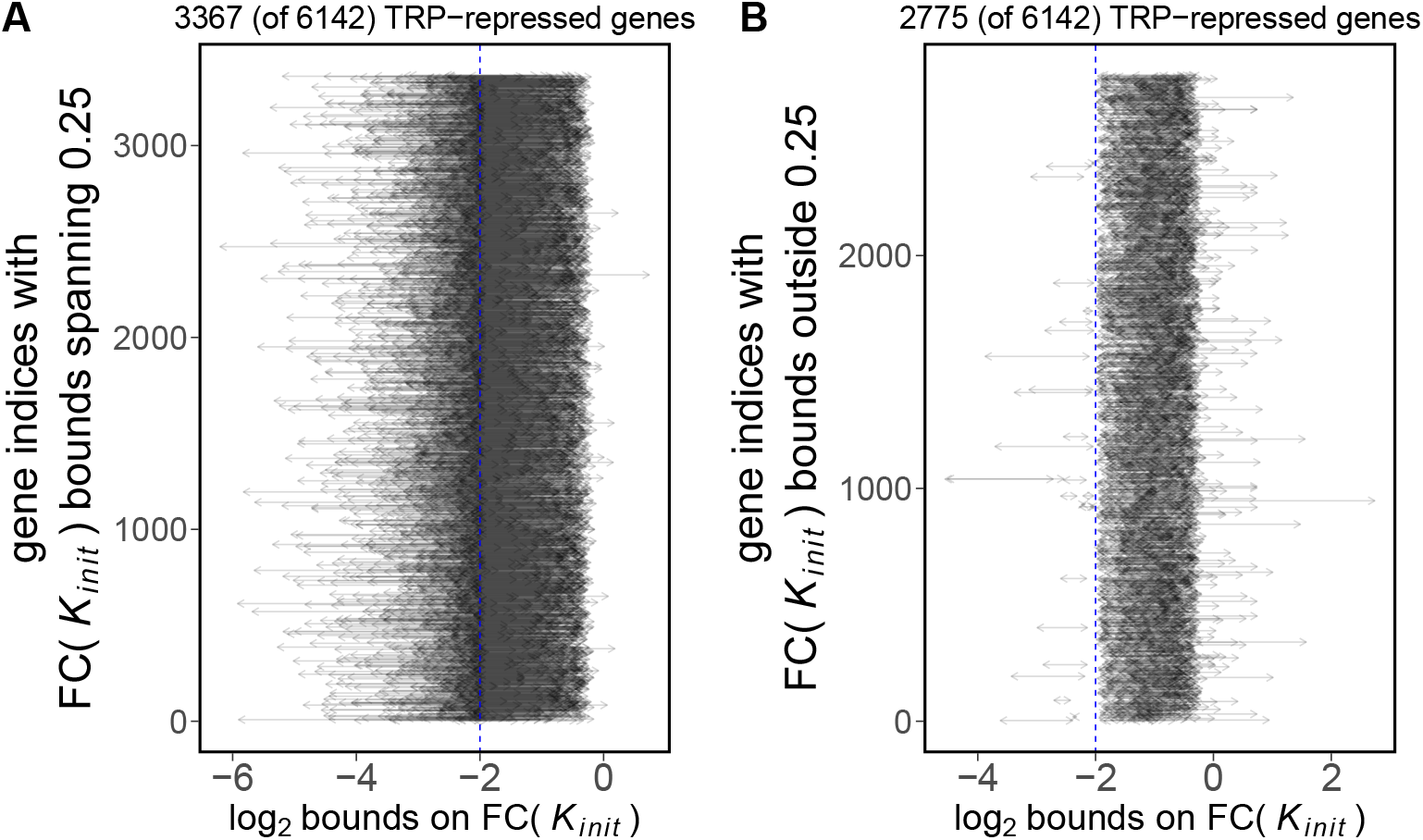
The model restrains the fold change initiation rates after triptolide inhibition for each genes within a calculated range. We consider repressed genes from a dataset that used 0.5 µM triptolide for 12.5 mins (11). A previous study reported a 0.25 fold change (dashed vertical line) in *K*_*init*_ upon treatment with 1µM triptolide at approximately 12 mins (55). We found 3326 (54.2% of 6131) TRP-repressed genes had bounds on changes in initiation rate spanning 0.25 (A). 2805 (45.7% of 6131) TRP-repressed genes had bounds on changes in initiation rate not spanning 0.25 (B). To be consistent with 75% decrease in initiation after triptolide treatment, we didn’t consider the 2805 genes in (B) while calculating ratio of premature termination to pause release (Eq. 17).

**Fig. S7.**
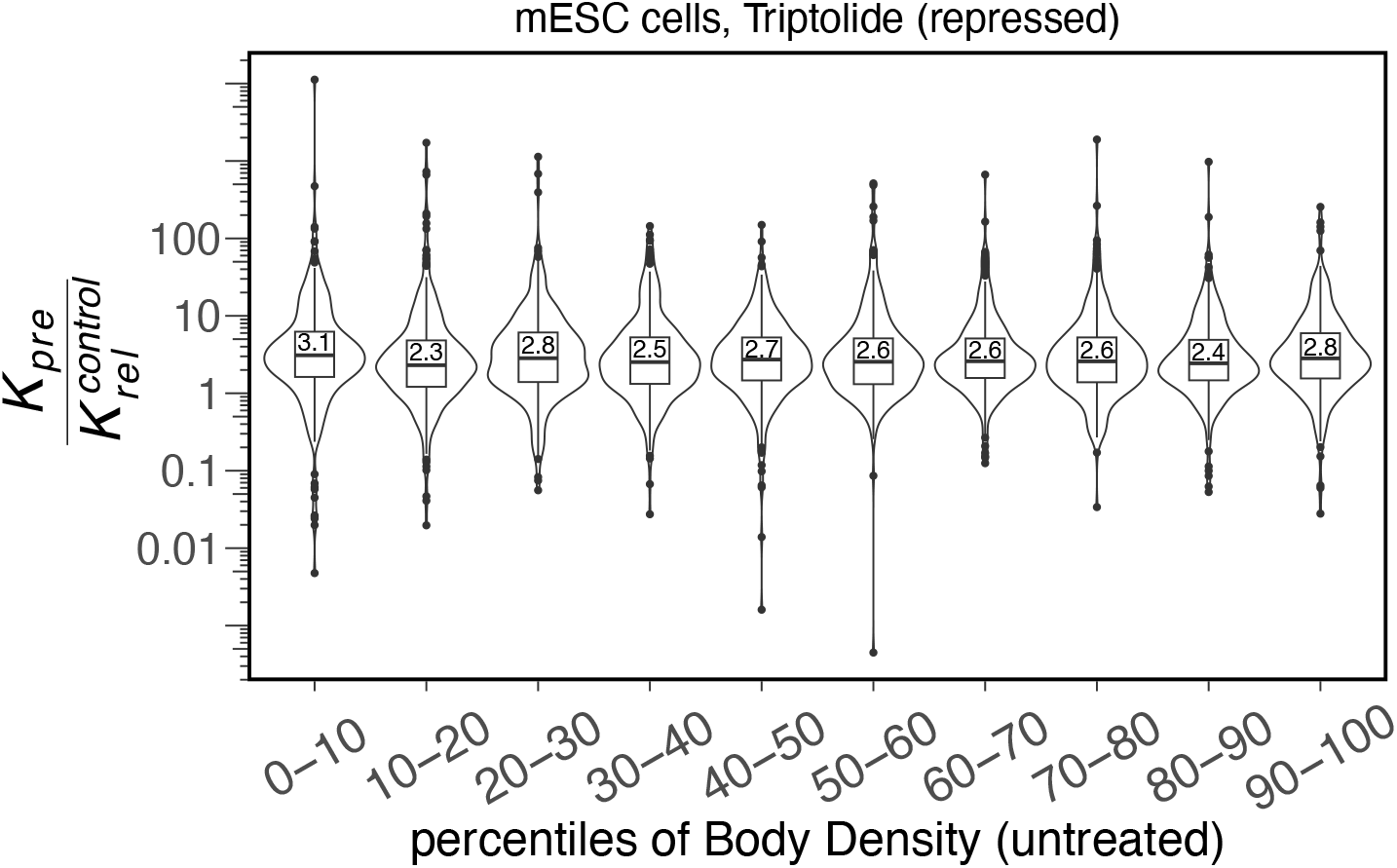
Premature termination is faster than pause release regardless of gene expression levels. We assumed that the repressed genes decrease initiation by ~75% and calculated ratio of premature termination and pause release rates for the indicated gene expression quantiles.

**Fig. S8.**
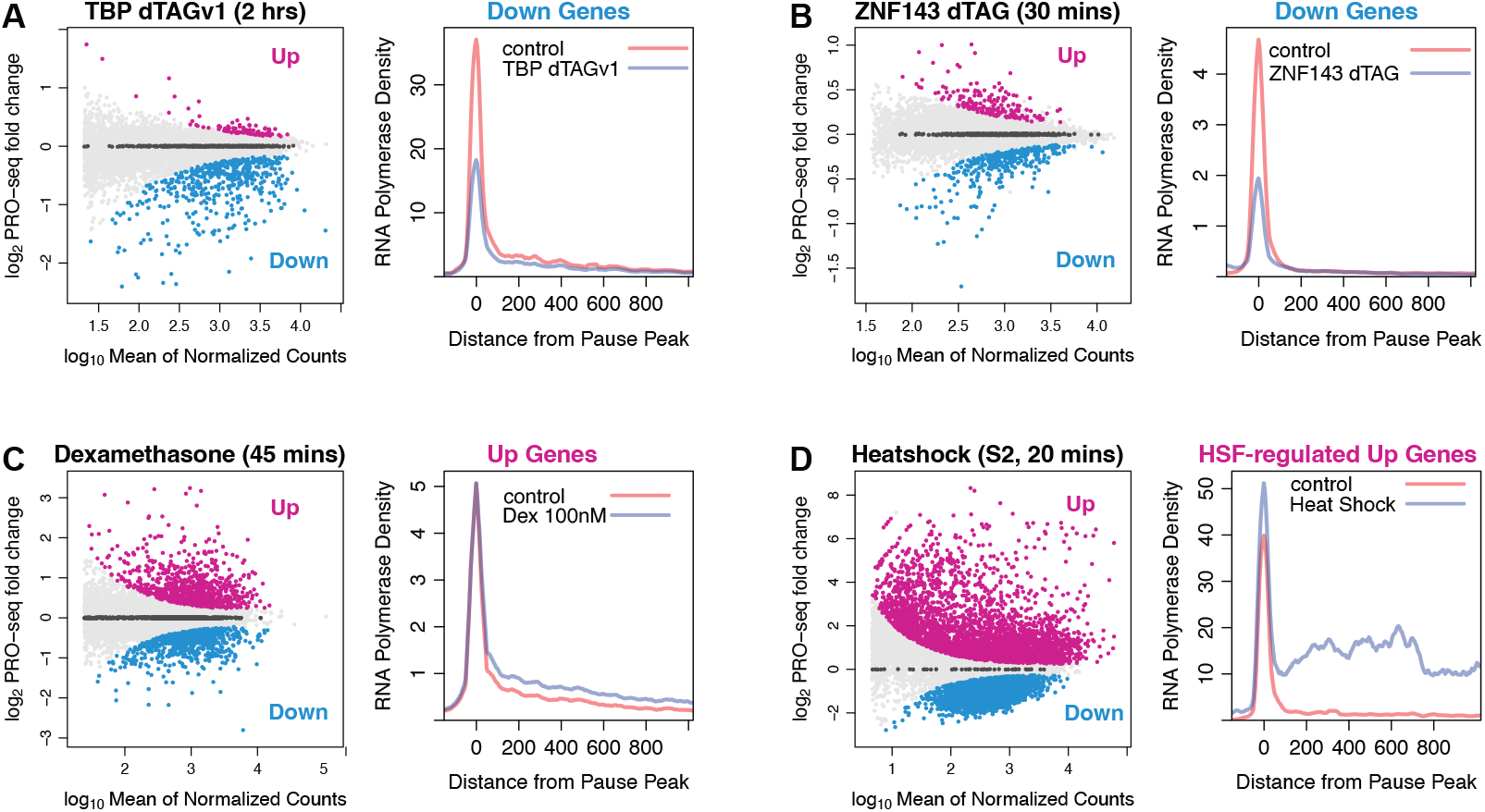
Acute treatments induce genomic changes in gene expression and RNA polymerase distribution. A) Degradation of TBP (43) led to 493 *Down* and 86 *Up* genes. The *Down* genes have reduced RNA polymerase density in pause region upon TBP degradation. B) Rapid degradation of ZNF143 (44) led to 358 *Down* genes and 181 *Up* genes. The *Down* genes decrease pause density. C) Dexamethasone treatment for 45 mins (45) resulted in 862 *Up* and 738 *Down* genes. The *Up* genes have increased RNA polymerase density in pause window and gene body upon dexamethasone treatment. D) Heat shock (HS) in *Drosophila S2* cells resulted in 3922 *Up* genes, although many of these genes are false positives due to read-through transcription (33). The genes that are confidently HSF-activated (Materials & Methods) increase RNA polymerase density in both the pause region and gene body.

**Fig. S9.**
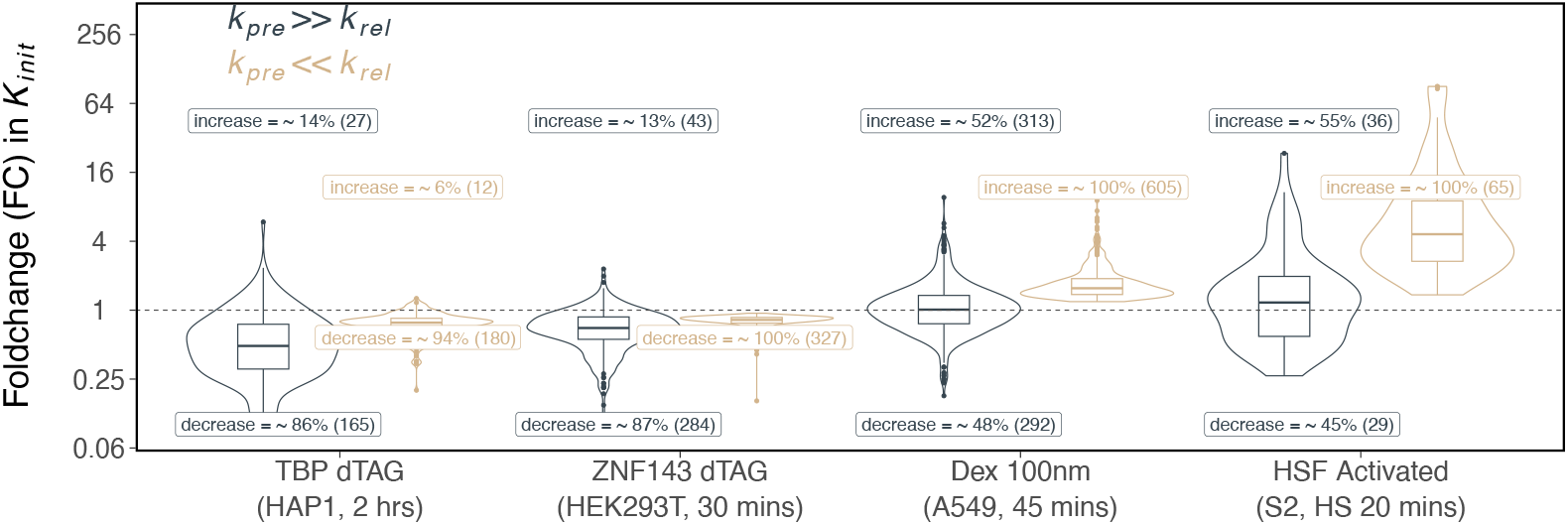
Changes in initiation rate for datasets used in this study vary substantially depending on the *k*_*rel*_ >> *k*_*pre*_ assumption. This figure reproduces the data from Figure 4 and includes changes in initiation rates for faster pause release as well (*k*_*rel*_ >> *k*_*pre*_).

**Fig. S10.**
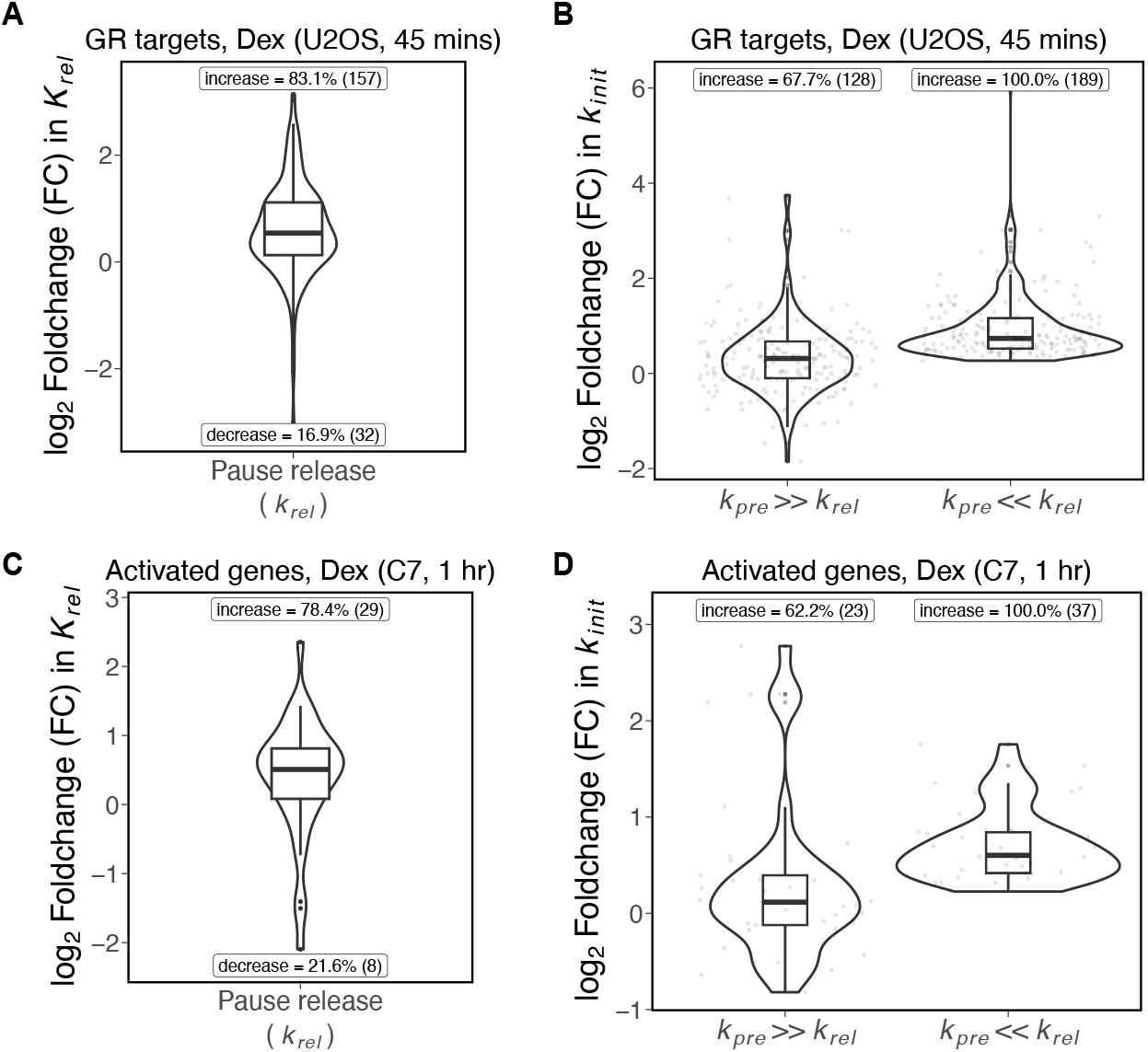
Dexamethasone induced GR binding leads to increase in pause release for activated genes in U2OS and C7 cells. We analyzed dexamethasone activated genes in U2OS cells (45) and C7 cells (46). A-B) We selected Dex-activated genes with TSS within 10kb of GR binding sites (81). 83.1% of activated genes had an increase in pause release, with moderate tendency of increase in initiation rate for 68% genes. C-D) We selected GR-target genes in C7 cells (46). 78.4% of activated genes had an increase in pause release with moderate increase in initiation rate for 62% genes.

**Fig. S11.**
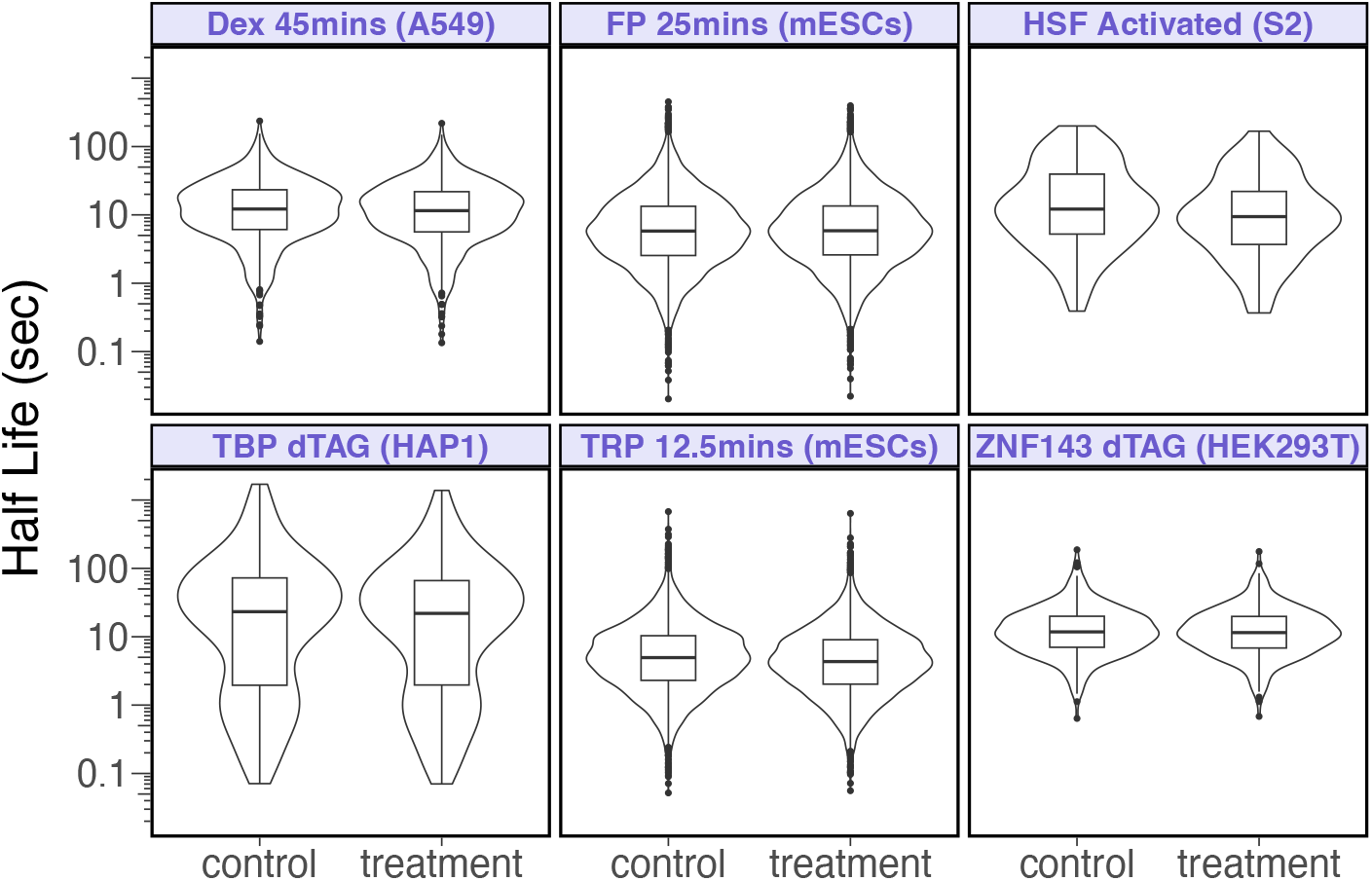
Turnover times of paused polymerases can become extremely rapid, reaching sub-second timescales at high premature termination rates. Violin plots illustrate the distribution of paused RNA polymerase half lives if premature termination rates at each gene promoter are ten times faster than pause release rates (i.e., *k*_*pre*_ = 10 × *k*_*rel*_). The median half-life of paused RNA polymerases across all datasets and treatments is 5.83 seconds, with an inter-decile range of 1.2 to 26 seconds.

**Fig. S12.**
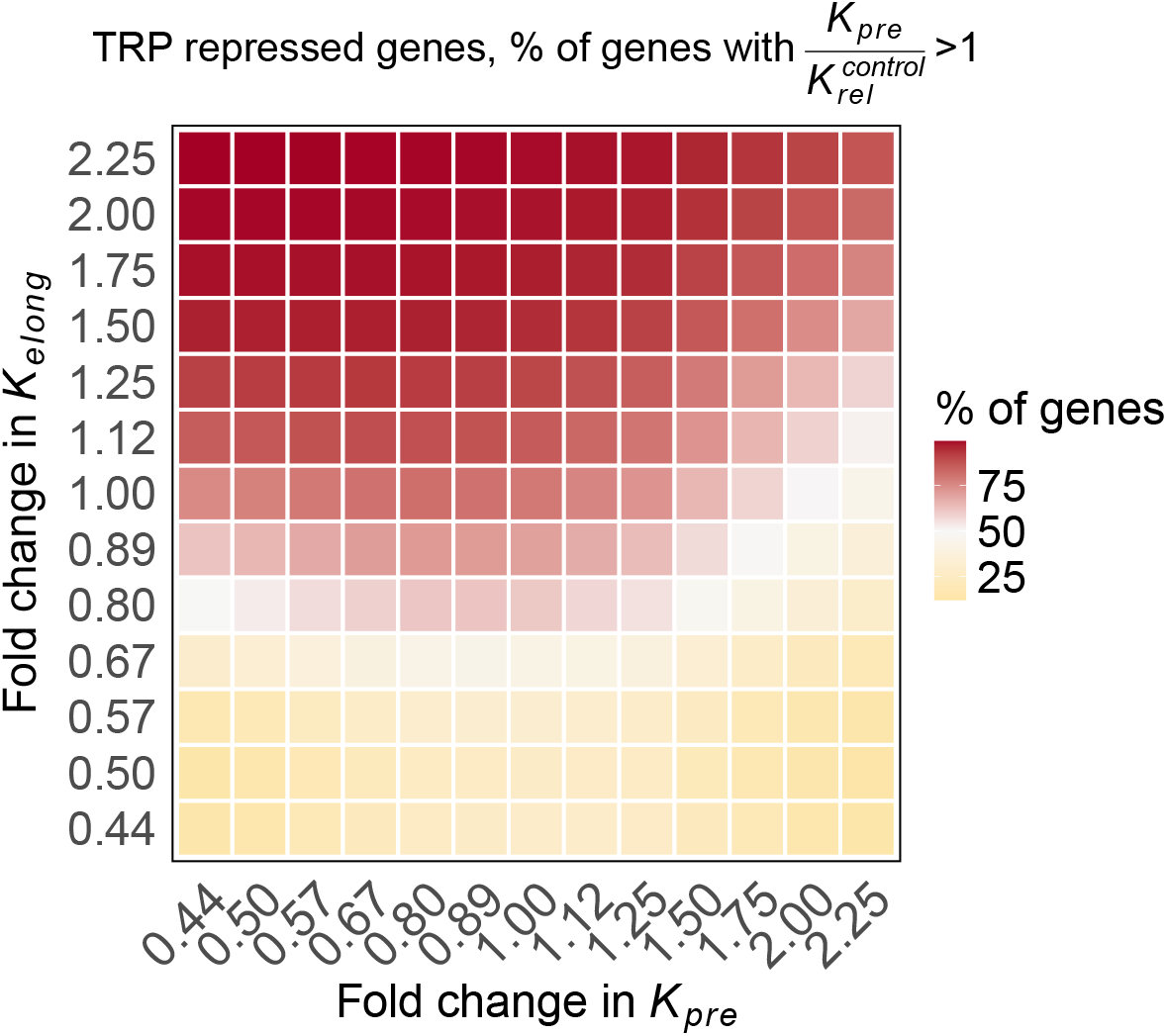
Premature termination is faster than pause release for most genes if triptolide doesn’t decrease elongation rate. For triptolide (TRP) repressed genes, we modeled each gene to have a 75% decrease in initiation. We calculated ratio of premature termination to pause release using (Eq. 16. To ensure the ratio in Eq. 16 remained positive, we selected genes that had 0.25 within the range defined by *FC*(*K*_*pre*_) × *FC*(*p*) and *FC*(*K*_*elong*_) × *FC*(*b*) for various values shown on the heatmap. White zones indicate the transition point where 50% of genes have faster premature termination.

**Fig. S13.**
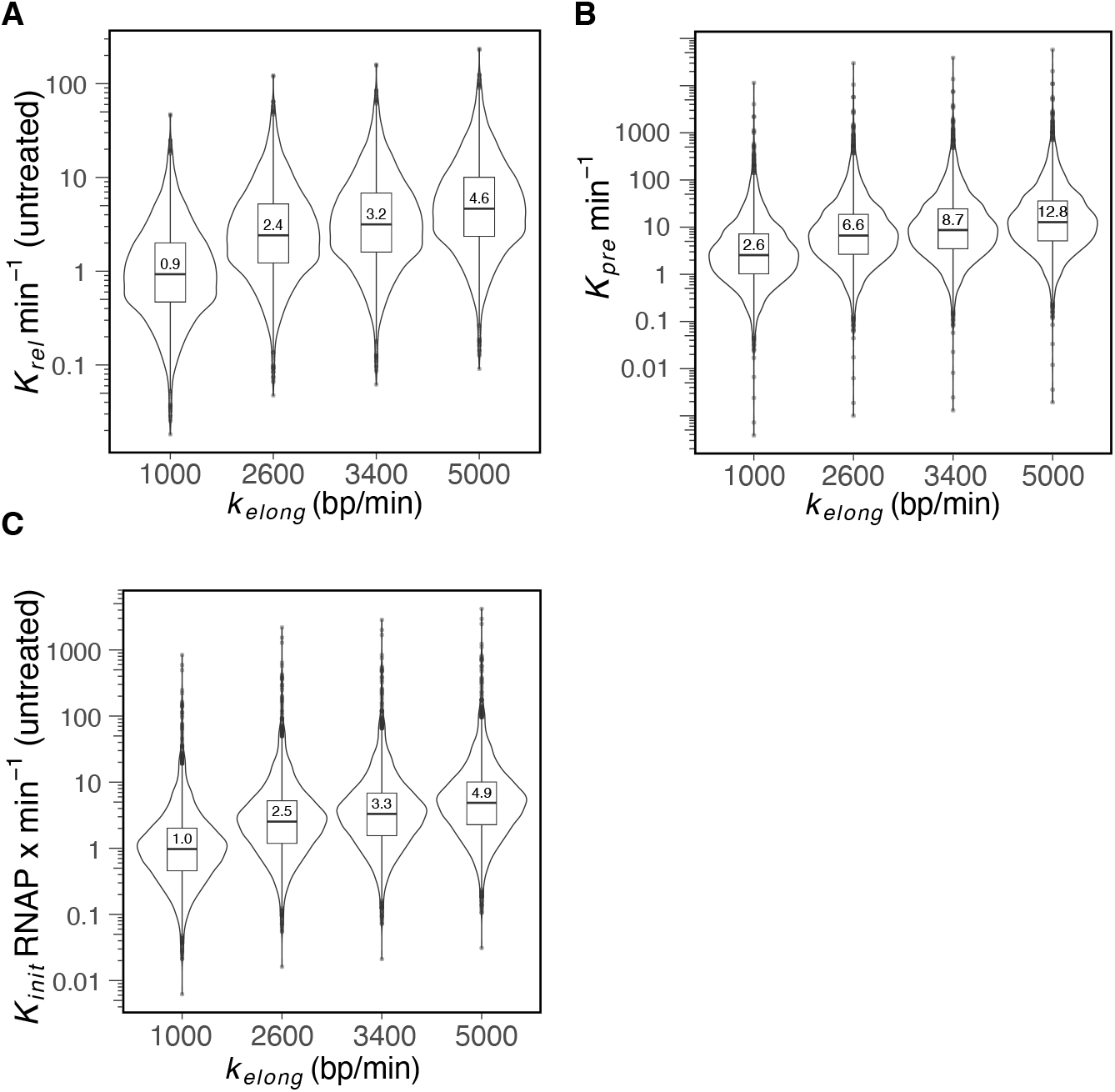
The rates of pause release, premature termination, and initiation are proportional to elongation rates. We varied elongation rates from 1–5 kb/min and considered TRP-repressed genes with bounds on changes in initiation rate spanning 0.25 (Fig. S6A). A) Under untreated conditions, pause release rates (*k*_*rel*_) had interdecile ranges of 0.24–4.2 events/min, 0.6–11 events/min, 0.81–14.2 events/min, and 1.2–21 events/min for elongation rates of 1, 2.6, 3.4, and 5 kb/min, respectively. Median values are indicated in the boxplots. B) We calculated premature termination rates (*k*_*pre*_) based on pause release using Eq. 17. The fold change in initiation rate (*k*_*init*_) was set to 0.25. The inter-decile ranges of premature termination rates were 0.5-20.4 events/min, 1.3-53 events/min, 1.7-69.2 events/min and 2.5-102 events/min for elongation rates of 1, 2.6, 3.4, and 5 kb/min. C) We derived initiation rates (*k*_*init*_) using values of premature termination and elongation rates (Eq. 14). The inter-decile ranges of initiation rates were 0.23-4.7 RNAP/min, 0.6-12.2 RNAP/min, 0.8-16 RNAP/min and 1.1-23.4 RNAP/min for 1, 2.6, 3.4, and 5 kb/min.

**Fig. S14.**
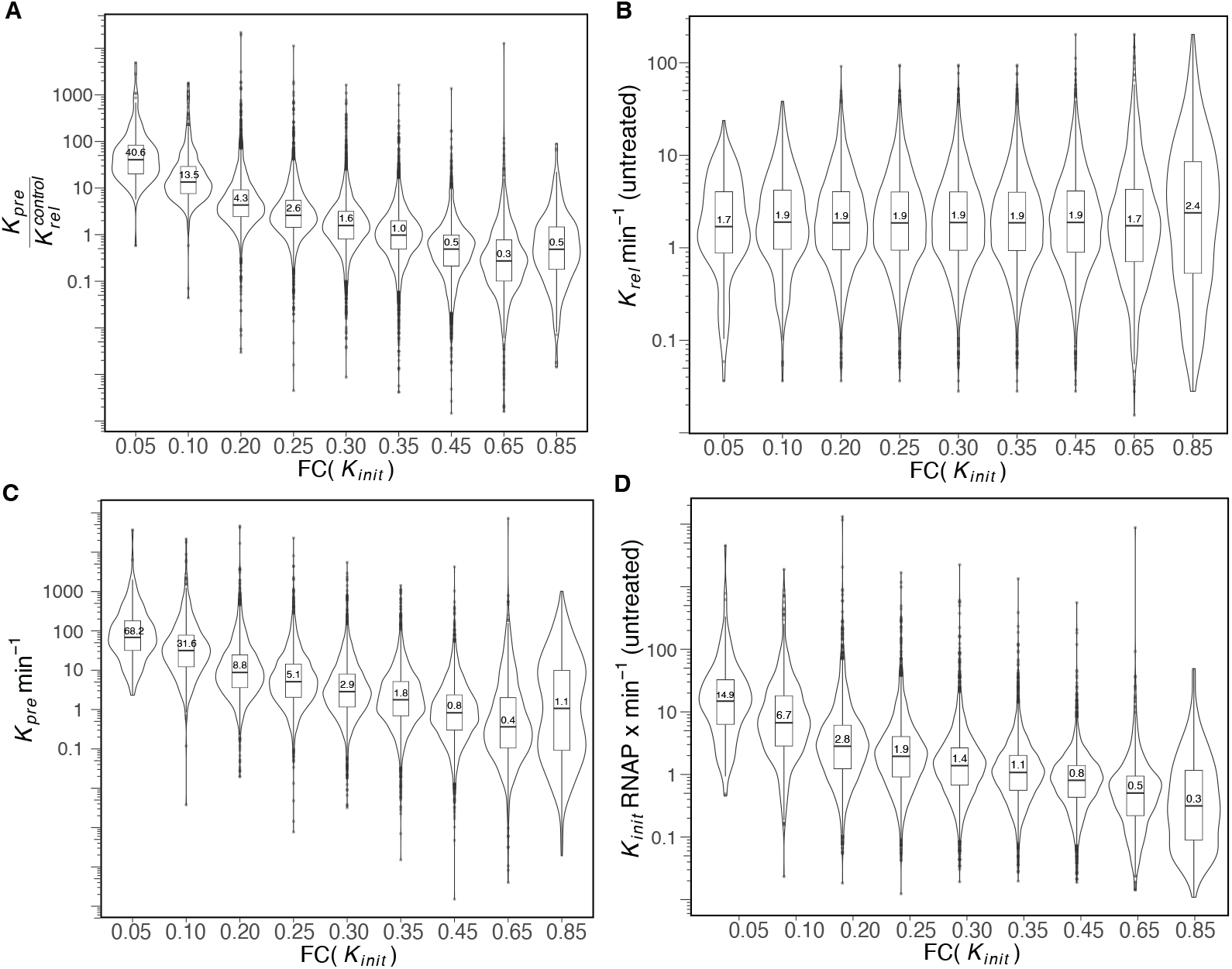
Parameter sensitivity analysis indicates that the premature termination to pause release rate ratio is >1 if triptolide reduces initiation rate to 0.35 times (or less) the control initiation rate. We varied the global decrease in *k*_*init*_ from 95% to 15%, corresponding to a fold change (FC) in *k*_*init*_ from 0.05 to 0.85 for repressed genes under TRP-inhibition (11). We considered the genes with bounds on fold-change in initiation rate (Eqs. 11–12) spanning the fold changes. We set the elongation rate at a representative value of 2000 bp/min. A) We calculated ratio of premature termination to pause release using Eq. 17. The inter-decile range at FC (*k*_*init*_) = 0.05 is 13.4-222, decreasing to inter-decile range of 0.4-7.2 at FC (*k*_*init*_) = 0.3, 0.21-4.4 at FC(*k*_*init*_) = 0.35 and 0.04-2.2 at FC(*k*_*init*_) = 0.65. B) Considering elongation rate of 2000 bp/min and set of genes with limits spanning the corresponding FC(*k*_*init*_), the pause release rates (Eq. 13) have an inter-decile range of 0.3-7.8 events/min at FC(*k*_*init*_) = 0.05, 0.5-8.3 events/min at FC(*k*_*init*_) = 0.35 and 0.3-11.3 events/min at FC(*k*_*init*_) = 0.65. C) We estimated premature termination (*k*_*pre*_) based on the ratio in (A). The inter-decile range of *k*_*pre*_ is 10.5-486 events/min at FC (*k*_*init*_) = 0.05, decreasing to 0.5 - 23 events/min at FC(*k*_*init*_) = 0.3, 0.03-11.8 events/min at FC(*k*_*init*_) = 0.65. D) We calculated initiation rate as indicated in Eq. 14. The inter-decile range changes from 2.1-85.2 RNAP/min at FC(*k*_*init*_) = 0.05 to 0.3 - 3.9 RNAP/min at FC (*k*_*init*_) = 0.35 and 0.09 to 2 RNAP/min at FC(*k*_*init*_) = 0.65.

**Fig. S15.**
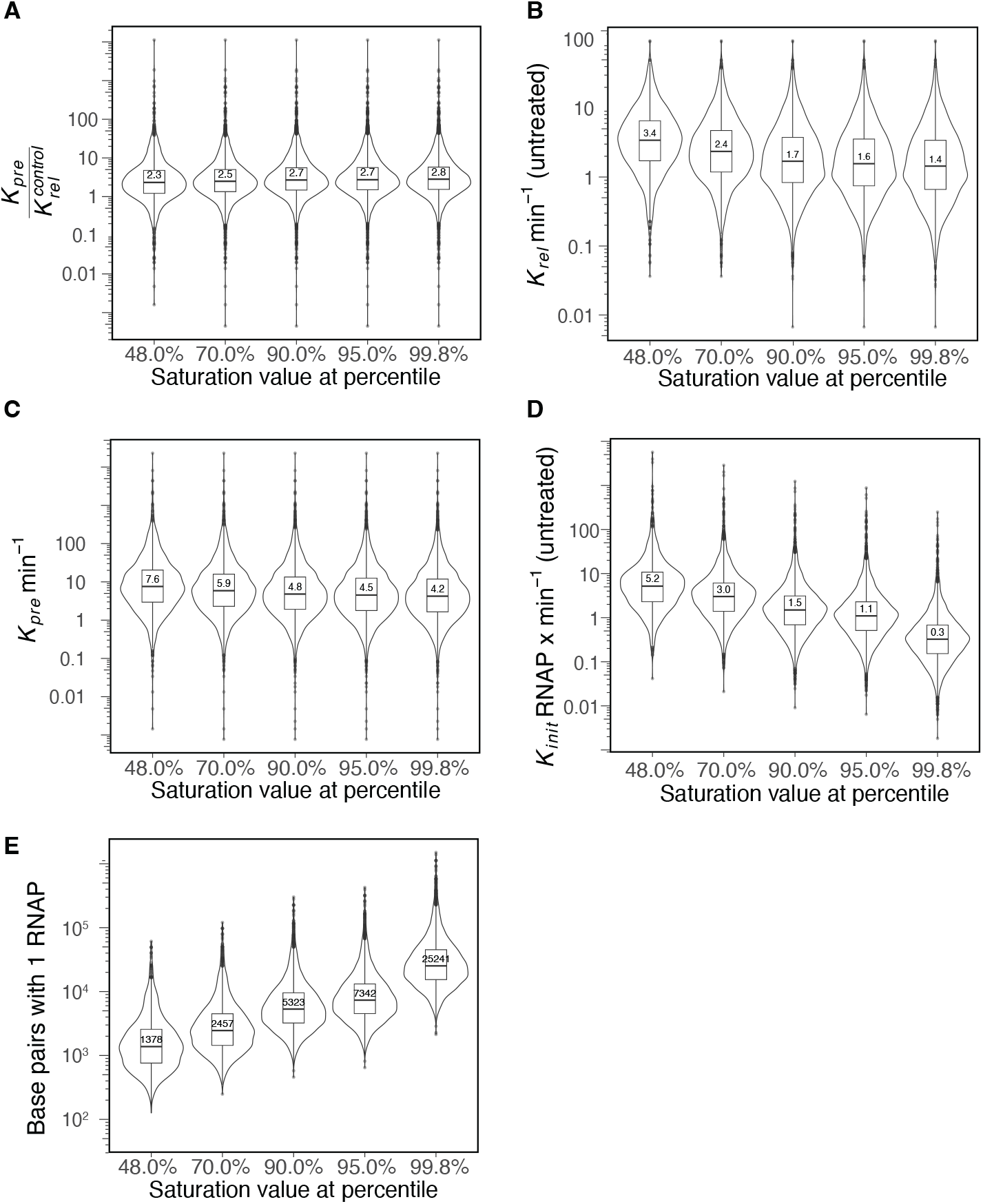
Increasing signal from a fully occupied pause region decreases rates of pause release, initiation and premature termination. We selected saturation values at different percentiles of ranked pause sums for repressed genes under TRP-inhibition (11). To achieve 25-fold dynamic range around the saturation value considered at 90th percentile of ranked pause sums (P = 169, 90th percentile), we varied saturation values from 34 (at 48th percentile) to 844 (at 99.8th percentile). A) We calculated ratios of premature termination and pause release using Eq. 17. We set the foldchange (FC) in initiation rate to 0.25 and elongation rate to 2000 bp/min. The inter-decile ranges of ratio is 0.6-12 at 48th percentile, 0.8-13.8 at 95th percentile. B) The pause release rates (Eq. 13) have an inter-decile range of 0.86-12 events/min at 48th percentile and 0.32-7.2 events/min at 99.8th percentile. C) The premature termination rates (Eq. 17) have an inter-decile range of 1.3-58.6 events/min at 48th percentile to 0.7-34 events/min at 99.8th percentile. D) The initiation rates (Eq. 14) have an inter-decile range of 1.1-24.7 polymerase/min at 48th percentile and 0.07-1.6 polymerase/min at 99.8th percentile. E) We calculated the density of RNA Polymerase on gene body using Eq. 15. The RNA polymerases were less densely spaced on gene body with increasing signal of a fully occupied pause region. The inter-decile ranges were 1 polymerase every 508 bp - 5.5 kb at 48th percentile which decreased to 1 polymerase every 2.2 kb - 18.4 kb at 90th percentile and to 1 polymerase every 10.2 kb - 87 kb at 99.8th percentile.

**Fig. S16.**
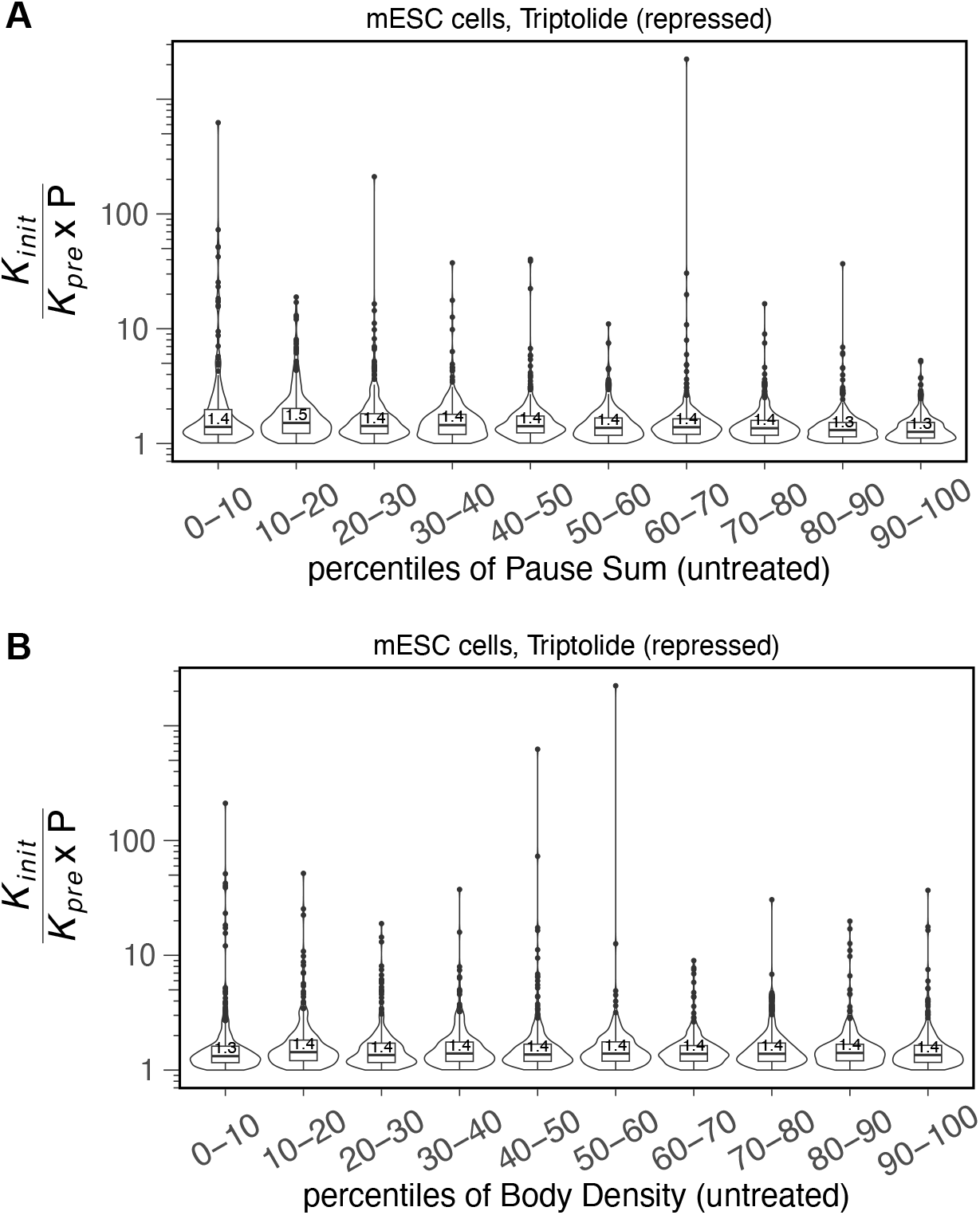
Most initiated RNA polymerases prematurely terminate. We categorized genes based on percentiles of paused polymerase density (A) and gene expression (B). We then estimated the ratio of initiation rate to effective premature termination (*k*_*pre*_ × *P*) under untreated conditions. The median values ranged from 1.3 to 1.5, with a distribution highly skewed towards 1. This suggests that the effective premature termination rate (*k*_*pre*_ × *P*) is approximately 67-77% of *k*_*init*_, regardless of paused polymerase density or gene expression levels.

**Table S1.**
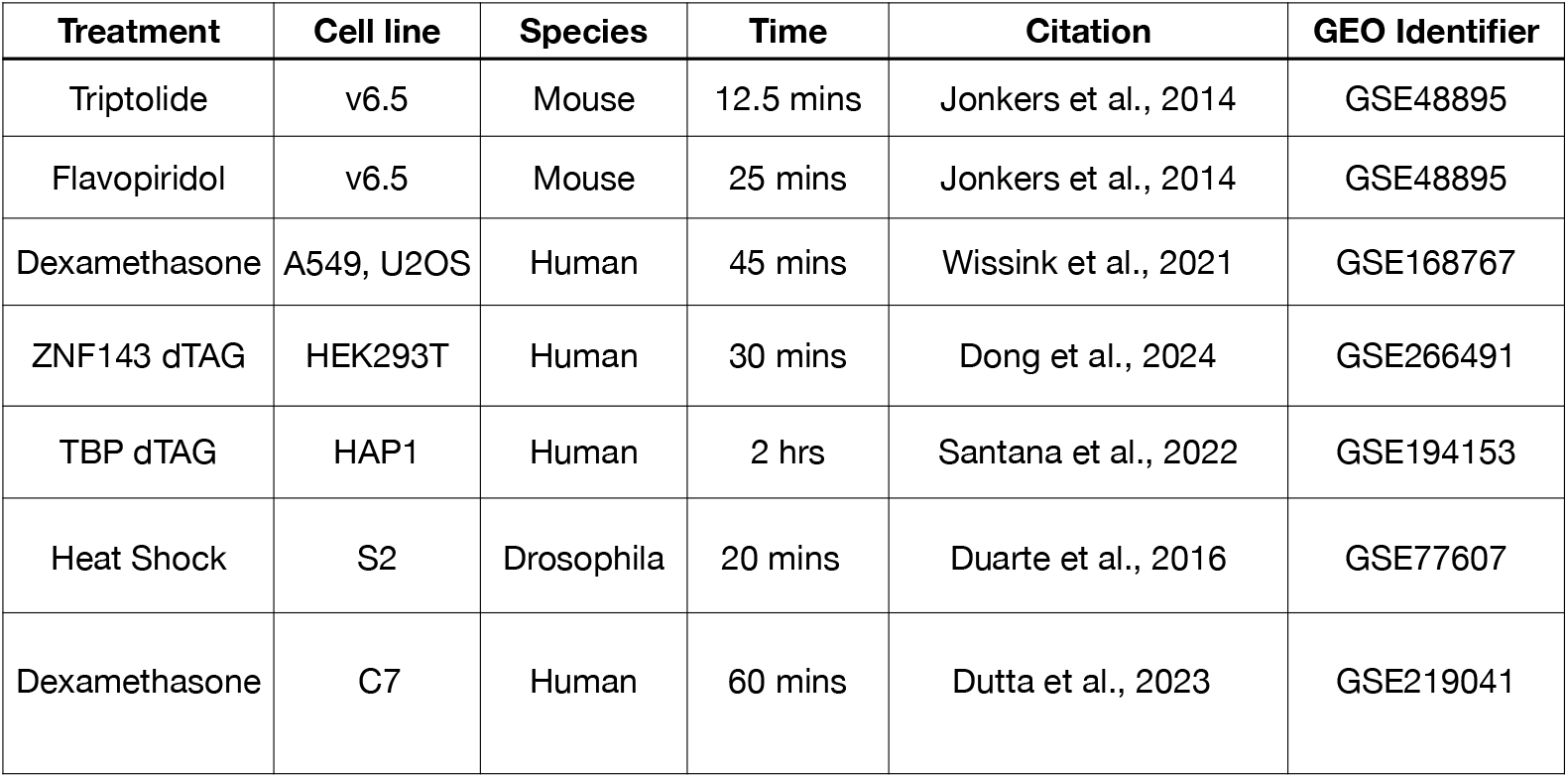
Gene Expression Omnibus (GEO) accession numbers for datasets used in this study.

**Table S2.** Multiple independent studies report faster premature termination rate compared to pause release.

| **Study** | **Premature Termination** | **Pause Release** |
| --- | --- | --- |
| Zimmer et al., 2021 | median 0.11 min<sup>-1</sup>; inner 90% range 0.027–0.23 min<sup>-1</sup> | median 0.027 min<sup>-1</sup>; inner 90% range 0.0015–0.31 min<sup>-1</sup> |
| Steurer et al., 2018 | Frequency: 12 out of 13 Pol2 terminate at promoter proximal pause region (~92.4%, Fig. 3B) | Frequency: 1 out of 13 Pol2 proceed into elongation (7.6%, Fig. 3B) |

**Table S3.** We report the median rates of the untreated samples from datasets with Unique Molecular Identifiers (UMIs). We complement the rate constants by including the effective rates, which refer to the number of RNA polymerase molecules per minute.

| **Treatment** | **Dataset** | **Pause release** | **Premature termination** | **Effective pause release** | **Effective premature termination** | **Initiation rate** |
| --- | --- | --- | --- | --- | --- | --- |
| Dexamethasone (A549) | Wissink et al. 2021 | 0.86/min | 2.24/min | 0.21RNAP/min | 0.55RNAP/min | 0.76RNAP/min |
| TBP dTAG (HAP9) | Santana et al., 2022 | 0.45/min | 1.2/min | 0.15RNAP/min | 0.38RNAP/min | 0.53RNAP/min |
| ZNF143 dTAG (HEK293T) | Dong et al., 2024 | 0.89/min | 2.32/min | 0.21RNAP/min | 0.55RNAP/min | 0.76RNAP/min |

## Notes

### Competing Interest Statement

The authors have declared no competing interest.

### Summary of Updates

The newer version has some additional analyses that reviewers requested, specifically an expansion of the sensistivity analysis.

